# Geometry-Constrained Coupling Drives Critical Dynamics Across Neural Manifolds

**DOI:** 10.1101/2025.09.07.674721

**Authors:** Richa Phogat, Anna Behler, Saurabh Sonkusare, James C. Pang, Nikitas Koussis, James A. Roberts, Jordan DeKraker, Sasha Dionisio, James M. Shine, Alex Fornito, P. A. Robinson, Michael Breakspear

**Affiliations:** School of Science, The University of Newcastle, Callaghan, New South Wales, Australia; QIMR Berghofer Medical Research Institute, Brisbane, Australia; School of Psychological Sciences, Turner Institute for Brain and Mental Health, and Monash Biomedical Imaging, Monash University, Clayton, Victoria, Australia; Mark Hughes Foundation Centre for Brain Cancer Research, College of Health, Medicine and Wellbeing, University of Newcastle, Callaghan, New South Wales, Australia; Brain Modelling Group, QIMR Berghofer, Herston, Brisbane, Queensland, Australia; Montreal Neurological Institute and Hospital, McGill University, Montreal, Canada; Neuroscience Center of Excellence, King Faisal Specialist Hospital and Research Centre, Riyadh, Saudi Arabia; School of Medical Sciences, Faculty of Medicine and Health, The University of Sydney, Sydney, New South Wales, Australia; School of Physics, University of Sydney, Camperdown, New South Wales, Australia; School of Medicine and Public Health, College of Health, Medicine and Wellbeing, University of Newcastle, Callaghan, NSW, Australia

## Abstract

Numerous physical systems evolve on curved manifolds whose geometry constrains their dynamics. The human brain provides a canonical example, with large-scale neural dynamics evolving on distinct anatomical surfaces such as the cortex and hippocampus. These surfaces support characteristic intrinsic dynamics shaped jointly by their geometry and internal feedback architecture. We capture both these influences in our geometry-constrained neural-field theory of cortical and hippocampal dynamics. In the brain, topographically organised reciprocal interactions couple these two systems, giving rise to collective cortico-hippocampal dynamics. Extending our theory to these collective dynamics therefore requires a coupling framework for fields evolving on geometrically distinct surfaces lacking a natural point-to-point correspondence. We achieve this by introducing a geometry-constrained coupling scheme that uses quasi-conformal mapping to place the two surfaces in a shared coordinate system, enabling spatially organised interactions between them. The resulting coupled system exhibits distinct dynamical regimes that capture characteristic signatures of recorded physiological and pathological cortico-hippocampal data. At moderate coupling, topographically linked regions show enhanced coordination while retaining their intrinsic cortical alpha (8-12 Hz) and hippocampal theta (4-8 Hz) rhythms. Stronger coupling, combined with increased hippocampal excitability, produces coherent low-frequency activity that recapitulates key features observed in human and rodent hippocampal seizures. Together, our results establish geometry-constrained coupling as an organising principle by which distinct neural manifolds reshape their intrinsic dynamics and undergo collective state transitions.

## I. INTRODUCTION

Spatially extended dynamics are constrained by the geometry of the manifolds on which they evolve. Curvature and topology can reorganise waves, patterns, and collective dynamics without altering the underlying local interaction rules [1–4]. The brain provides a particularly rich realisation of this general problem. Geometry shapes neural activity within individual anatomical structures [5], but the dynamical consequences of coupling across distinct neural manifolds remain poorly understood. This problem is central to the analyses and understanding of whole-field neurophysiological and functional neuroimaging data, where observed activity is shaped by both local dynamics and the coupling between major brain structures. The same challenge extends to pathological activity such as epileptic seizures, which typically involve interactions between multiple systems and circuits in the brain [6]. The cortico-hippocampal system provides a particularly important test case for this problem [7– The cortex and hippocampus are both spatially extended neural manifolds, but they differ markedly in geometry, scale, and anatomical organisation. The cortex forms a large, folded sheet, whereas the hippocampus is smaller, elongated, and curved along its longitudinal axis. Their coordination, therefore, poses a specific dynamical problem of integrating activity across geometrically distinct structures. These integrated dynamics underlie memory encoding, consolidation and retrieval [7–1 spatial navigation [16– and contextual processing [21]. A theory of cortico-hippocampal dynamics must therefore account for the intrinsic activity supported by each manifold and the emergent dynamics generated by spatially structured interactions between them.

A mechanistic account of this coupling must begin with an understanding of the anatomical pathways through which the cortico-hippocampal interactions occur. The hippocampus has widespread, reciprocal connections with neocortical regions, with many connections passing via the entorhinal cortex [22–26]. These connections are topographically organised, with specific cortical areas projecting to distinct regions of the hippocampus and vice versa [25, 27, 28]. This bidirectional flow of information creates a dynamic loop, enabling the hippocampus to act as a hub for coordinating and linking distributed cortical activity [26, 29]. Thus, the relationship between the hippocampus and cortex is characterised by a spatially organised, functionally adaptive, and reciprocally interconnected architecture that underpins their shared role in cognition [30].

These anatomical pathways specify the possible routes of interaction, but they do not by themselves determine how activity propagates, synchronises, or reorganises across coupled neural structures. Capturing these dynamics requires a biophysical modelling framework that can represent population-level activity over continuous anatomical geometries while remaining computationally tractable. Prevailing macroscopic mean-field approaches for simulating brain dynamics are commonly formulated either via neural mass models (NMMs) or neural field theories (NFTs) [31]. NMMs approximate brain regions as discrete nodes connected via empirical structural networks [32–36], and have been used extensively to study cortical and subcortical interactions, including wave-like activity [37, 38]. However, their node-based formulation limits their ability to capture activity constrained by the geometry of extended neural structures, such as the folds and undulations of the cortex or the elongated curvature of the hippocampus. This limitation is particularly relevant in light of recent work showing that physical geometry strongly constrains neural activity [5].

NFT provides a complementary spatially continuous formulation by modelling population-level activity as a field evolving over anatomical space [39–44]. In the NFT framework, neural activity propagates across spatially extended tissue, allowing surface geometry, local neighbourhood structure, and physiological parameters to shape the resulting dynamics [44–48]. NFT can therefore provide valuable insight into the biophysical processes that generate large-scale cortical rhythms [43, 44, 48].

Despite this broader capability, existing NFT formulations have focused primarily on the corticothalamic system, where reciprocal loops between the cortex and thalamus explain the emergence of various neocortical rhythms, including cortical alpha activity [43, 44, 48– 50]. Recent advances in hippocampal surface reconstruction now allow the folded archicortical anatomy of the hippocampus to be represented as a continuous surface, analogous to surface-based representations of the cortex [51, 52]. Together with accumulating evidence for wave-like activity propagating across the hippocampus [53–55], these developments motivate a spatially continuous description of hippocampal dynamics. The hippocampus exhibits a characteristic theta rhythm in the 3–8 Hz range, shaped in part by reciprocal interactions with the medial septum, which provides rhythmic excitatory and inhibitory input [56, 57]. The corticothalamic and hippocampo-septal systems therefore share a common organising principle in which spatially extended neural activity is shaped by reciprocal feedback with a subcortical structure.

Extending NFT to the hippocampo-septal system therefore provides a second spatially continuous field, evolving on a geometrically distinct anatomical manifold. Because cortical and hippocampal activity is strongly interdependent in the brain, these two fields cannot be understood solely through their individual dynamics. This raises the broader question of coupling continuous active fields on geometrically distinct surfaces while respecting the topographic organisation of the anatomical interactions between them.

We address both aspects of this problem by starting with separate neural field models of the corticothalamic and hippocampo-septal systems, capturing their intrinsic cortical and hippocampal dynamics. We then introduce a geometrically constrained coupling between the cortical and hippocampal surfaces using quasi-conformal mapping [58]. This mapping enables interactions between the two fields while preserving local neighbourhood relationships. By systematically increasing the coupling strength, we show that inter-manifold interactions reshape intrinsic cortical and hippocampal rhythms and induce topographically organised synchrony. When combined with increased hippocampal excitability, the same coupling mechanism drives a transition to seizure-like spectral and synchronisation dynamics that recapitulate key features of human and rodent intracranial EEG (iEEG) recordings. Put together, our results suggest that the geometry-constrained coupling between these two neural manifolds acts as an organising principle for their collective dynamics during both health and pathology.

## II. RESULTS

### A. Overview

We first formulate and simulate separate neural field models of the corticothalamic and hippocampo-septal systems. In these models, cortical and hippocampal activity evolves on population-averaged anatomical surfaces embedded in three-dimensional space. Distributed feedback through the thalamus and septum shapes the intrinsic rhythms of the cortex and the hippocampus, respectively. The dynamics of each field are governed by coupled integro-differential equations, which we solve on the corresponding anatomical surface using a modal decomposition in geometric eigenmodes. This decomposition provides a low-dimensional representation of the fields, enabling computationally efficient simulations.

To investigate how interactions between the cortex and the hippocampus shape their collective dynamics, we next introduce spatially structured coupling between their anatomical surfaces. We use quasi-conformal mapping [58] to project the geometrically distinct cortical and hippocampal surfaces into a shared two-dimensional Euclidean domain. By approximately preserving local neighbourhood relationships, this transformation enables a spatially structured and proximity-based coupling between the two surfaces. We then systematically increase the coupling strength to determine how cortico-hippocampal interactions reorganise the emergent dynamics. The resulting spectral predictions are compared with cortical and hippocampal intracranial electroen-cephalographic (iEEG) recordings from humans and rodents.

### B. Neural Field Theory of Corticothalamic Activity

NFT describes population-level neural activity as continuous fields evolving over anatomically structured space. In the corticothalamic neural field model, 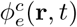 denotes the cortical excitatory activity at location **r** and time *t* as it propagates across the cortical surface. This activity is maintained by recurrent intracortical interactions and delayed thalamocortical feedback, and evolves according to a damped wave equation [5, 42, 59] as follows,

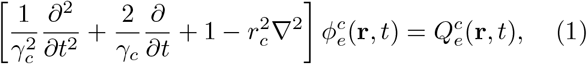

where *c* indicates those variables and parameters associated with the corticothalamic subsystem and the subscript *e* denotes the cortical excitatory population. The spatial operator ∇^2^ describes the lateral propagation of activity on the folded cortical surface. The parameter *r*_*c*_ denotes the characteristic range of cortical axonal projections, which follows the well-documented exponential distance rule (i.e., the probability and weight of connections between two locations decrease as an exponential function of their physical separation [60]). The damping rate (*γ*_*c*_) controls the temporal attenuation of the propagating activity. The source term 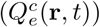 represents the mean firing rate of the cortical excitatory population and drives the evolution of 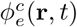.

The corticothalamic NFT system comprises four neural populations: local excitatory (*e*) and inhibitory (*i* ) cortical populations, along with the reticular (*r* ) and relay nuclei (*s*) of the thalamus (Fig. 1a). The total presynaptic input to a population is written as 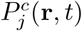, where *j* ∈ {*e, r, s*}. These inputs are constructed by summing the field activity arriving from all populations that project to *a*, weighted by the corresponding connection strengths 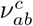. The cortical excitatory field, 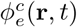, propagates across the cortical sheet as well as providing input to thalamic populations, after a corticothalamic conduction delay *τ*_*c*_. In contrast, thalamic activity does not propagate laterally as a field within the thalamus. The presynaptic cortical excitatory input 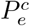 receives contributions from local cortical excitation, local cortical inhibition, and delayed relay activity from the thalamus. The reticular nuclei of the thalamus receive input 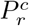 from the relay thalamic nuclei and delayed cortical excitation. Lastly, the input to the relay nuclei 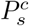 consists of delayed cortical excitation, reticular inhibition, and nonspecific subcortical drive with stochastic fluctuations. The total incoming activity to each population is therefore given by,

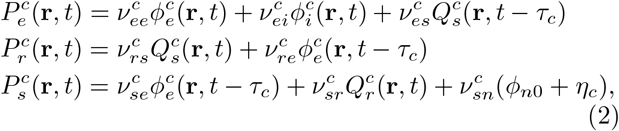

where 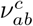 denotes the synaptic connectivity strength to population *a* from population *b*, as indicated by the subscripts. The corticothalamic loop incorporates the conduction delay (*τ*_*c*_) between cortex and thalamus. *ϕ*_*n*0_ represents a baseline nonspecific input and *η*_*c*_ is a stochastic input term. Due to their predominantly local connections, the inhibitory cortical field 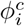 is assumed to locally follow the excitatory cortical field 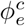 without supporting spatial propagation. The expression for the inhibitory population 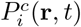 is analogous to 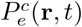 and is omitted for brevity.

**FIG. 1.**
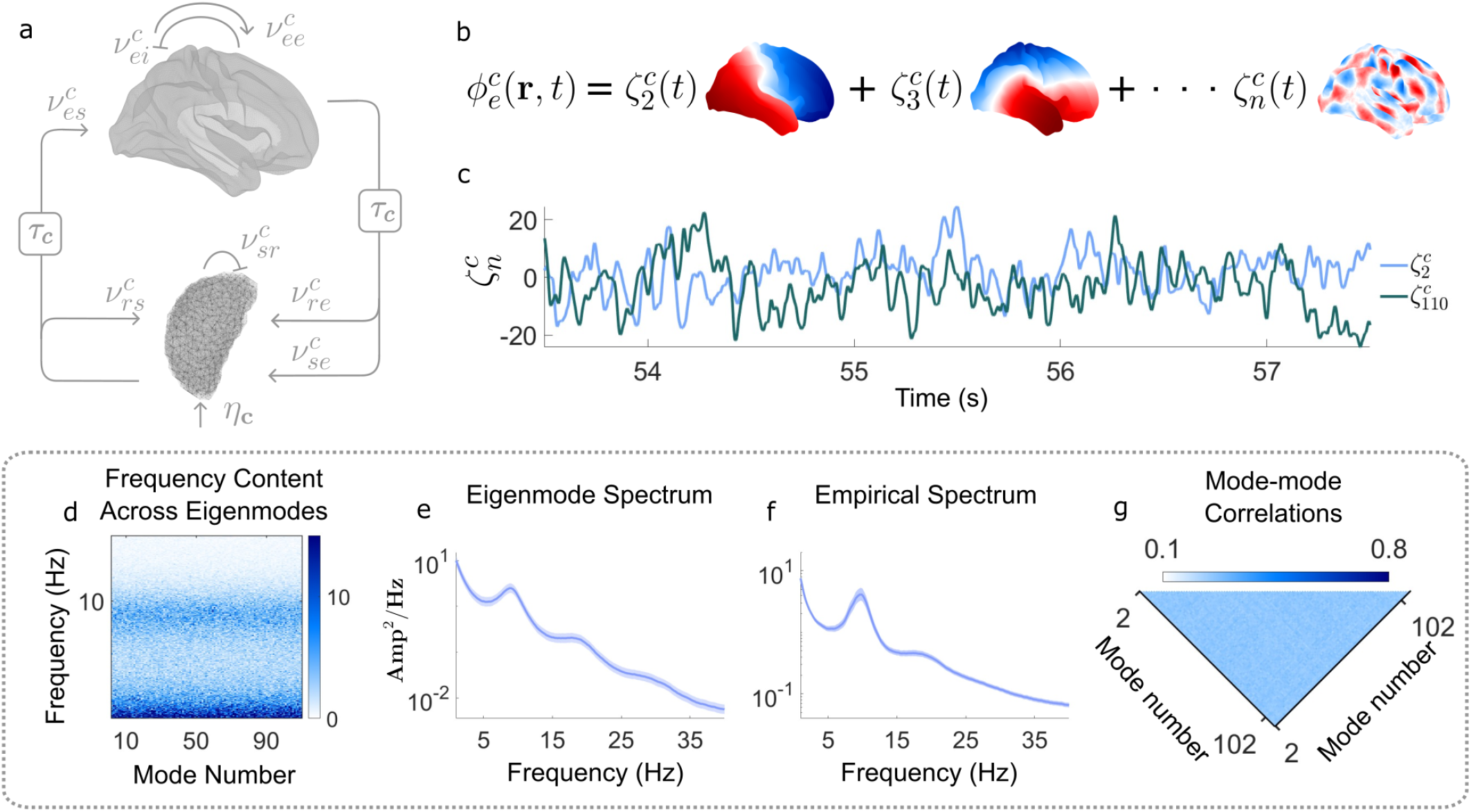
Corticothalamic system dynamics. **a**, Schematic of the corticothalamic model where the letters denote the interactions between the excitatory (*e*), inhibitory (*i* ) populations of the cortex and the reticular (*r* ) and relay (*s*) nuclei of the thalamus. **b**, Cortical activity is decomposed into geometric eigenmodes, ensuring a compact spatio-temporal representation. **c**, Representative eigenmode amplitude time series of the 2nd and 110th eigenmodes illustrate the evolution of cortical activity at different spatial scales. **d**, Power spectra of the individual eigenmode time series, displayed as a heatmap, reveal spectral profiles of different eigenmodes. **e**, Average PSD of the cortical eigenmode time series is shown in dark blue. **f**, Average cortical PSD computed from resting-state EEG data collected from 64 EEG electrodes across 111 participants. The light blue shading in both **e** and **f** indicates the smoothed 95% confidence interval of the mean PSD. **g**, Dynamic synchrony measure (DSM), computed as the windowed maximum lagged cross-correlation between eigenmode time series, reflects uniformly weak correlations across spatial scales.

The local transmembrane potential 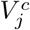 is then generated from the aggregate presynaptic input 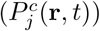, for each population, filtered through synaptic and dendritic processes. The governing equations for 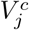 are expressed as,

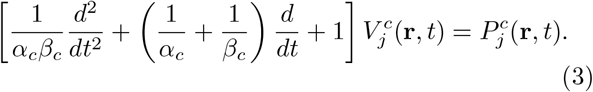

The impulse response of this synaptodendritic filter follows a biexponential postsynaptic kernel, with a rapid rise and slower decay governed by the rate constants *α*_*c*_ and *β*_*c*_.

The neural field equations are then completed by converting the resulting local mean transmembrane potentials into the mean firing rate 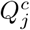 through a sigmoidal activation function,

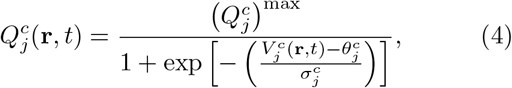

where 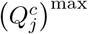 represents the maximum firing rate of each of the four neural populations *j* ∈ {*e, i, r, s*}. The parameters 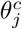 and 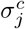 denote the mean firing threshold and its standard deviation, respectively. The resulting mean firing rates 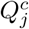 then become the source term for the damped wave equation, closing the system of field equations.

Using this framework, corticothalamic dynamics were simulated with previously published physiologically derived parameters (Table I, Methods), incorporating empirically-informed wave speeds, damping rates, firing rates, neural gains, and synaptodendritic rate constants [43, 44, 48, 59]. Numerical integration of the damped wave equation 1 was performed via modal decomposition, in which cortical activity at each integration step is projected onto geometric eigenmodes of the cortex (Fig. 1b and Methods section A). This representation expresses cortical activity as a weighted sum of cortical eigenmodes with time-varying amplitudes, providing a compact description of the system’s spatiotemporal dynamics. In doing so, it also incorporates the influential role of cortical geometry on large-scale neuronal dynamics [5, 50, 61]. Lower-order eigenmodes correspond to large-scale patterns of activity, whereas higher-order modes capture shorter wavelengths (Fig. 1b).

**TABLE I.** Cortical model parameters.

| Parameter | Description | Value |
|---|---|---|
| *v*<sub>*c*</sub> | Axonal conduction velocity | 11.6 m s<sup>−1</sup> |
| *r*<sub>*c*</sub> | Characteristic spatial range | 0.1 m |
| *γ*<sub>*c*</sub> = *v*<sub>*c*</sub>/*r*<sub>*c*</sub> | Axonal damping rate | 116 s<sup>−1</sup> |
| *Q*<sub>*cj*</sub><sup>max</sup> | Maximum firing rate | 340 s<sup>−1</sup> |
| *θ*<sub>*j*</sub><sup>*c*</sup> | Firing threshold | 12.92 × 10<sup>−3</sup> V |
| *σ*<sub>*j*</sub><sup>*c*</sup> | Threshold std. deviation | 3.4 × 10<sup>−3</sup> V |
| *α*<sub>*c*</sub> | Inverse decay time | 83.33 s<sup>−1</sup> |
| *β*<sub>*c*</sub> | Inverse rise time (*β*<sub>*c*</sub>/*α*<sub>*c*</sub> = 9.23) | 769.14 s<sup>−1</sup> |
| *τ*<sub>*c*</sub> | Corticothalamic delay (one-way) | 0.04 s |
| *ν*<sub>*ee*</sub><sup>*c*</sup> | Cortical excitation *e* → *e* | 3.03 × 10<sup>−3</sup> V·s |
| *ν*<sub>*ei*</sub><sup>*c*</sup> | Cortical inhibition *i* → *e* | −6.00 × 10<sup>−3</sup> V·s |
| *ν*<sub>*es*</sub><sup>*c*</sup> | Thalamic relay *s* → cortical excitation *e* | 2.06 × 10<sup>−3</sup> V·s |
| *ν*<sub>*se*</sub><sup>*c*</sup> | Cortical excitation *e* → thalamic relay *s* | 1.65 × 10<sup>−3</sup> V·s |
| *ν*<sub>*sr*</sub><sup>*c*</sup> | Thalamic reticular *r* → relay *s* | −0.83 × 10<sup>−3</sup> V·s |
| *ν*<sub>*sn*</sub><sup>*c*</sup> | Nonspecific input *n* → relay *s* | 0.98 × 10<sup>−3</sup> V·s |
| *ν*<sub>*re*</sub><sup>*c*</sup> | Cortical excitation *e* → reticular *r* | 0.33 × 10<sup>−3</sup> V·s |
| *ν*<sub>*rs*</sub><sup>*c*</sup> | Thalamic relay *s* → reticular *r* | 0.03 × 10<sup>−3</sup> V·s |
| *ϕ*<sub>*n*0</sub> | Baseline nonspecific input | 1 |

This eigenmode-based implementation of the corti-cothalamic NF model does not require computationally intensive simulation of cortical activity at every surface vertex. Instead, the dynamics of the cortical sheet are projected onto a compact basis of cortical eigenmodes, allowing large-scale spatiotemporal activity to be represented by a low-dimensional set of mode amplitudes. The temporal evolution of two representative cortical eigenmodes illustrates how the model captures the spatiotemporal dynamics shaped by cortical geometry (Fig. 1c). The resulting eigenmode time series exhibit an apparent alpha-band (≈10 Hz) activity, captured by the peak in their power spectral density (PSD), considered individually (Fig. 1d) and when averaged across eigenmodes (Fig. 1e). Alpha rhythms in the 8-12 Hz frequency range are a robust feature of resting-state cortical activity [33, 45]. For comparison, the empirical PSD computed from 64 scalp EEG electrodes and averaged across 111 participants from the OpenNeuro SRM resting-state EEG dataset [62] is shown in Fig. 1f. Simulated and empirical data clearly share the prominent alpha-band peak, with a smaller beta peak (≈ 20Hz) followed by a featureless spectral drop off at higher frequencies. Windowed maximum lagged correlations across all cortical eigenmodes (Fig. 1g) reveal that longer versus finer wavelength activity evolve with minimal mutual influence, with diffuse, weak cross-scale synchrony. Localised spatial correlations arising from relatively weak global synchronisation are hallmarks of nonpathological cortical dynamics [63, 64]. Hence, the eigenmode-based simulation retains the key spectral features of cortical activity [43, 44], while providing an efficient representation of its spatiotemporal dynamics with the activity of over 29,000 vertices efficiently simulated by just 110 cortical eigenmodes.

### C. Neural Field Theory of Hippocampal Activity

Accumulating evidence for the influence of the intrinsic geometric organization of the hippocampus on its activity [7, 16, 53, 54, 65] makes NFT a strong candidate for understanding hippocampal dynamics. Here we extend the NFT framework to the hippocampo-septal system and compare the predicted field 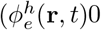 to its characteristic empirical spectra. In particular, we focus on hippocampal theta rhythms, which are spatiotemporally organized and strongly modulated by the septum [53, 56, 57, 66]. To maintain notational consistency, the NFT equations for the hippocampo-septal system adopt the same mathematical structure as the corticothalamic system with the exception that variables and parameters associated with the hippocampo-septal system are distinguished by using *h* instead of *c*.

Similar to the corticothalamic system, the evolution of the hippocampal excitatory field 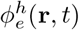 is governed by a damped wave equation,

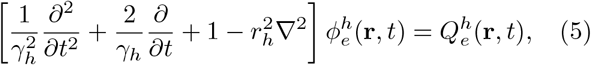

where the parameters *γ*_*h*_ and *r*_*h*_ represent the temporal damping rate and characteristic spatial range of hippocampal excitatory neural populations. Comparable to the corticothalamic system, the hippocampus and medial septum also form a feedback loop that plays a pivotal role in shaping the spatiotemporal dynamics of the hippocampal theta rhythm [56, 67, 68]. The time delay in this loop is given by *τ*_*h*_ (Table II, Methods section). The remaining equations governing source terms 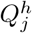 and transmembrane potentials 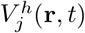 follow the same mathematical structure as the corticothalamic system [Equations (2), (3), and (4)], with parameters adjusted for the hippocamposeptal system. The subscripts *e* and *i* in the hippocampo-septal system denote excitatory and inhibitory populations within the hippocampus, while *r* and *s* refer to inhibitory and excitatory populations in the medial septum (Fig. 2a). Throughout the manuscript, hippocampal dynamics are shown in red and cortical dynamics in blue across all spectral and correlation visualizations.

**TABLE II.**
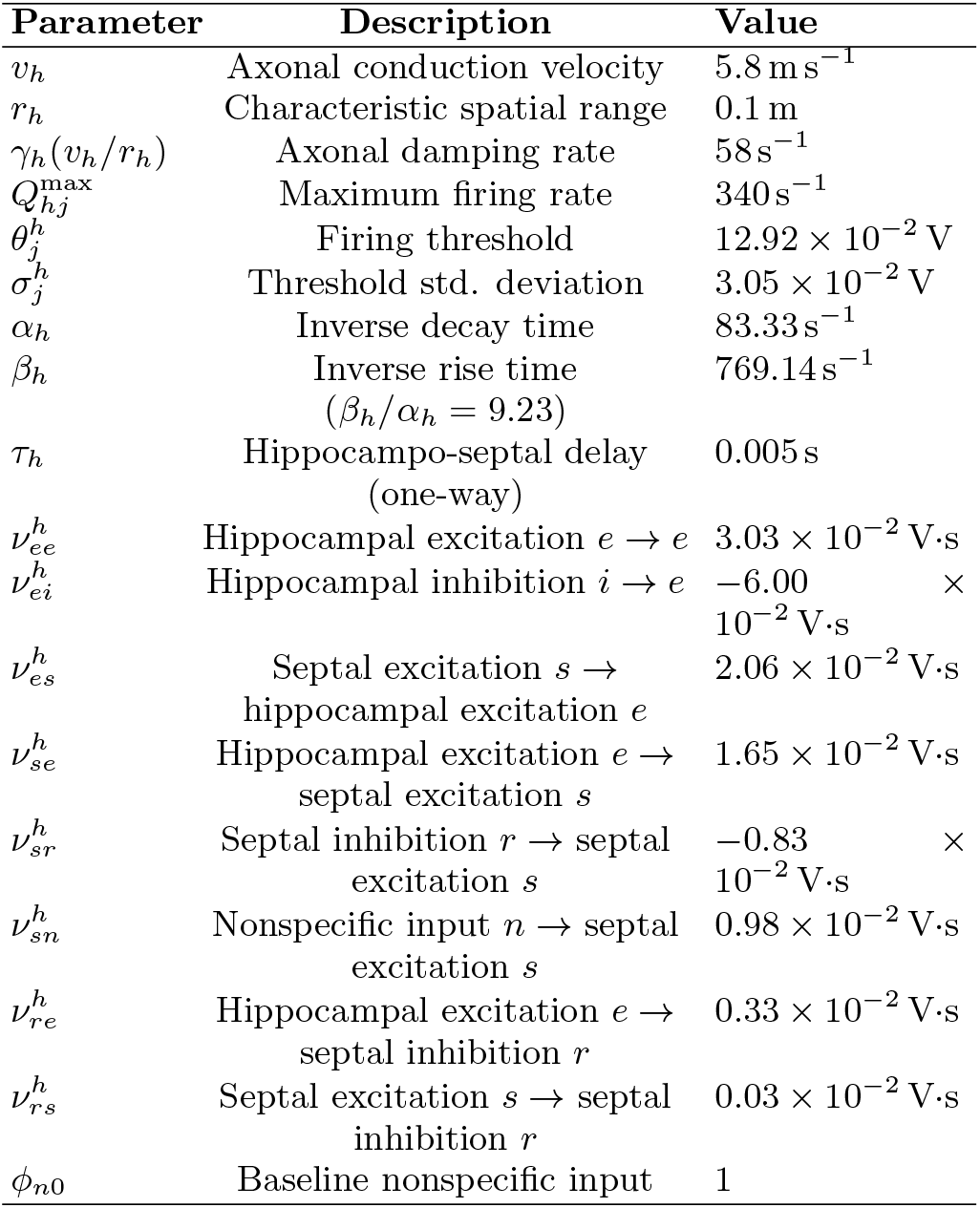
Hippocampal model parameters.

| Parameter | Description | Value |
|---|---|---|
| *v*<sub>*h*</sub> | Axonal conduction velocity | 5.8 m s<sup>−1</sup> |
| *r*<sub>*h*</sub> | Characteristic spatial range | 0.1 m |
| γ<sub>*h*</sub>(*v*<sub>*h*</sub>/*r*<sub>*h*</sub>) | Axonal damping rate | 58 s<sup>−1</sup> |
| *Q*<sub>*hj*</sub><sup>max</sup> | Maximum firing rate | 340 s<sup>−1</sup> |
| θ<sub>*j*</sub><sup>*h*</sup> | Firing threshold | 12.92 × 10<sup>−2</sup> V |
| σ<sub>*j*</sub><sup>*h*</sup> | Threshold std. deviation | 3.05 × 10<sup>−2</sup> V |
| α<sub>*h*</sub> | Inverse decay time | 83.33 s<sup>−1</sup> |
| β<sub>*h*</sub> | Inverse rise time (β<sub>*h*</sub>/α<sub>*h*</sub> = 9.23) | 769.14 s<sup>−1</sup> |
| τ<sub>*h*</sub> | Hippocampo-septal delay (one-way) | 0.005 s |
| ν<sub>*ee*</sub><sup>*h*</sup> | Hippocampal excitation *e* → *e* | 3.03 × 10<sup>−2</sup> V·s |
| ν<sub>*ei*</sub><sup>*h*</sup> | Hippocampal inhibition *i* → *e* | −6.00 × 10<sup>−2</sup> V·s |
| ν<sub>*es*</sub><sup>*h*</sup> | Septal excitation *s* → hippocampal excitation *e* | 2.06 × 10<sup>−2</sup> V·s |
| ν<sub>*se*</sub><sup>*h*</sup> | Hippocampal excitation *e* → septal excitation *s* | 1.65 × 10<sup>−2</sup> V·s |
| ν<sub>*sr*</sub><sup>*h*</sup> | Septal inhibition *r* → septal excitation *s* | −0.83 × 10<sup>−2</sup> V·s |
| ν<sub>*sn*</sub><sup>*h*</sup> | Nonspecific input *n* → septal excitation *s* | 0.98 × 10<sup>−2</sup> V·s |
| ν<sub>*re*</sub><sup>*h*</sup> | Hippocampal excitation *e* → septal inhibition *r* | 0.33 × 10<sup>−2</sup> V·s |
| ν<sub>*rs*</sub><sup>*h*</sup> | Septal excitation *s* → septal inhibition *r* | 0.03 × 10<sup>−2</sup> V·s |
| ϕ<sub>*n*0</sub> | Baseline nonspecific input | 1 |

**FIG. 2.**
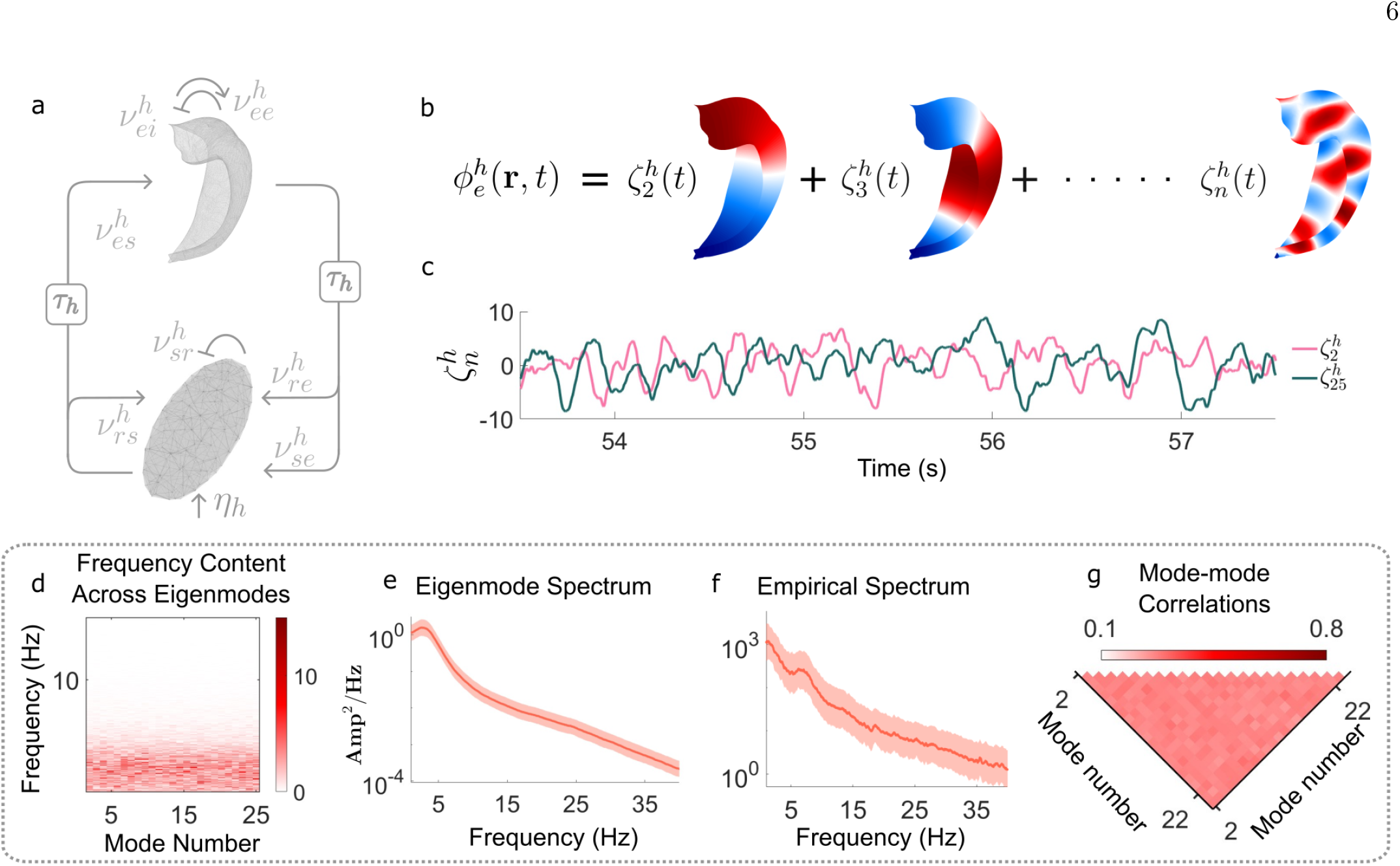
Hippocampo-septal system dynamics. **a**, Schematic of the hippocampo-septal model showing reciprocal connections between the hippocampus and the septum. **b**, Hippocampal activity is decomposed into eigenmodes, providing a compact representation of its spatiotemporal dynamics. **c**, Time series of the 2nd and 25th eigenmodes illustrate the evolution of hippocampal activity at different spatial scales. **d**, Frequency content across eigenmodes, visualized as a heatmap, highlights the spectral profile of each eigenmode. **e**, Average PSD across the simulated hippocampal eigenmode time series also reveals a peak in the theta-band (3-8 Hz). **f**, Average PSD from hippocampal electrodes in one subject is used as a representative example of human hippocampal theta activity. Dark lines indicate the mean PSD, and shaded regions indicate smoothed 95% confidence interval of the mean PSD in both **e** and **f. g**, Dynamic synchrony measure, quantified using a windowed maximum lagged cross-correlation between non-zero eigenmode time series, indicates weak correlations across hippocampal eigenmodes. The rationales for parameter selection and scaling in the hippocampo-septal NFT are described in the Methods section.

Hippocampal geometry is incorporated using a population-averaged surface mesh [52] of the hippocampus, from which geometric eigenmodes are derived. Adapting the same mode-based approach used for the cortex, the first 25 hippocampal eigenmodes are used for numerical simulations, capturing spatiotemporal patterns down to ≈ 2–3 mm resolution. Simulating this hippocampo-septal system within the NFT formalism yields robust theta-band activity (≈3–8 Hz). This is evident in both the representative hippocampal eigen-mode amplitude time series (Fig. 2c) and the PSD of the eigenmode time series (Fig. 2d and 2e). To compare the simulated hippocampo-septal spectrum with empirical human hippocampal activity, we computed an empirical hippocampal PSD from intracranial EEG recordings in the OpenNeuro dataset [69]. PSD from Hippocampal electrodes in subject 5 was used as a representative example of the spectral content of human hippocampal theta activity (Fig. 2f). As with the cortex, correlations between different hippocampal eigenmodes are diffuse and weak to moderate in strength (Fig. 2g) implying limited interactions across different spatial scales during healthy, non-pathological acquisitions.

Cortical alpha and hippocampal theta are considered canonical rhythms of these systems [7, 56]. Here we find that NFT models of these systems, considered separately, yield these physiologically realistic rhythms via anatomically grounded feedback loops and intrinsic dynamics.

### D. Cortico-Hippocampal Coupling

Cortical and hippocampal systems do not evolve independently but coordinate their dynamics via extensive reciprocal interactions [70]. Cortico-hippocampal interactions are organised along broad anatomical and functional gradients, particularly along the longitudinal axis of the hippocampal formation [23, 27, 28]. Modelling these interactions requires an empirically motivated rule for transferring spatially extended activity between two surfaces while preserving their topographic organisation (Fig. 3a). We implement this rule as a delayed, bidirectional, geometry-constrained coupling between the cortical and hippocampal fields, denoted by *C*_*ch*_ and *C*_*hc*_.

**FIG. 3.**
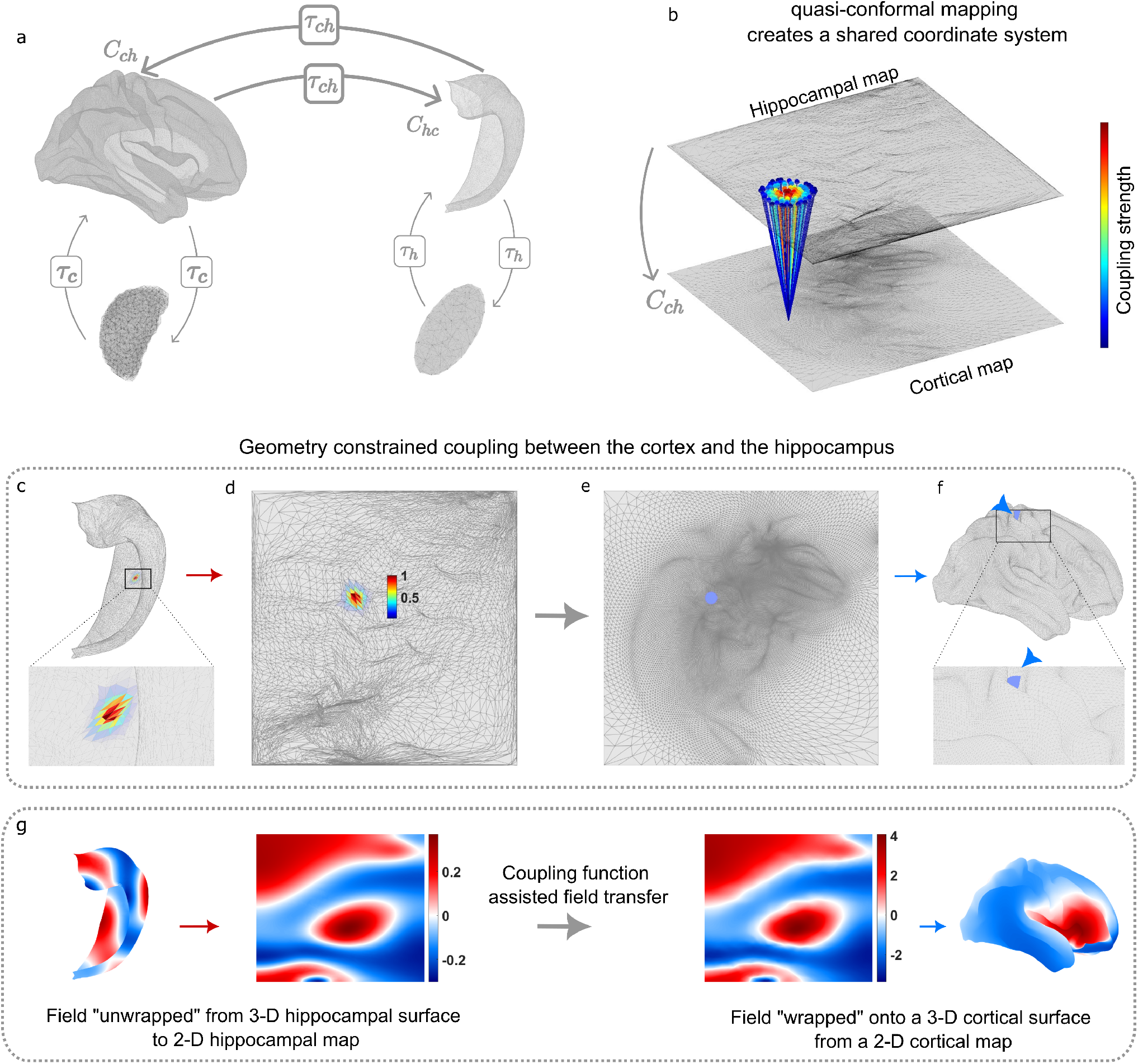
Coupled cortico-hippocampal system. **a**, The bidirectional connectivity between the cortex and hippocampus is represented by *C*_*ch*_ and *C*_*hc*_. **b**, Visualization of the hippocampus → cortex coupling enabling smooth field transfer between surfaces via quasi-conformal mapping. **c**–**f**, Detailed illustration of the mechanism underlying hippocampus → cortex field transfer. Both cortical and hippocampal surfaces are mapped onto a common 2-D domain via a quasi-conformal transformation (**b, d** and **e**, respectively). The target cortical vertex (**f** ) and its location on the 2-D cortical map (**e**). The hippocampal region (**d**) projecting to the target cortical vertex is computed via the coupling function 10, 11, which centres the coupling kernel *w*_*ch*_(**r** − **r**^*′*^) on the hippocampal vertex closest to the target cortical vertex in the 2-D coordinate space. The colour gradient in this region represents the coupling strength, peaking at the centre (red) and exponentially decaying outward. **g**, Illustration of the field propagation from the hippocampal surface to the cortical surface performed with the proposed coupling scheme. The same methodology can be implemented in the opposite direction (cortex to hippocampus) by swapping source and target maps.

Spatial continuity assumptions implicit in NFT require a coupling framework that preserves local neighbourhood relationships so that nearby locations on one surface interact with nearby regions on the other (Fig. 3b). Since the cortex and hippocampus form geometrically distinct manifolds, these relationships cannot be defined directly from their native coordinates. We hence apply quasi-conformal transformations to map both surfaces onto a common two-dimensional Euclidean domain (Fig. 3c–f). The resulting maps are then aligned along their anterior–posterior axes, encoding the empirical observation that anterior cortical regions interact preferentially with anterior hippocampal regions, and posterior cortical regions with posterior hippocampal regions, following core anatomical principles [23, 27, 28]. The cortical and hippocampal locations occupying the same or nearby positions in this shared domain are then mutually coupled.

Because neural fields represent population activity over spatially extended regions, coupling cannot be restricted to individual pairs of corresponding vertices. Instead, the input to each target vertex is obtained by combining activity over a neighbourhood on the source surface. To illustrate this construction, consider a target vertex on the cortical surface, shown by the blue point in Fig. 3e and 3f. Its position in the shared two-dimensional domain identifies the nearest corresponding hippocampal vertex, which defines the centre of our coupling kernel (Fig. 3b and 3d). The cortical target then receives input from the hippocampal region surrounding this centre in the shared Euclidean space. Activity at the central hippocampal vertex contributes most strongly to the coupling, while contributions from surrounding vertices decrease exponentially as a function of their distance from the kernel centre (Fig. 3c,d). These weighted contributions are integrated and transferred to the target cortical vertex (Fig. 3e,f).

As the target vertex moves across the cortical map, the corresponding kernel centre moves smoothly across the hippocampal map (Supplementary Materials Fig. S1–S3 and supplementary video S1). Neighbouring cortical vertices therefore receive input from neighbouring hippocampal regions, preserving local spatial organisation and avoiding abrupt changes or tears in the transferred field (Fig. 3g). Reversing the source and target surfaces defines the analogous cortex-to-hippocampus coupling.

The uncoupled wave equations of cortical and hippocampal fields can be written compactly as,

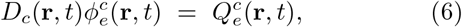

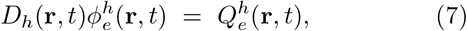

where *D*_*c*_ and *D*_*h*_ are the cortical and hippocampal differential operators introduced in Equations 1 and 5, respectively. These neural field equations are extended by incorporating spatially structured inter-surface interactions *W*_*ch*_(**r, r**′, *t, t*′) and *W*_*hc*_(**r, r**′, *t, t*′) in the source terms of Equations 1 and 5, yielding the full evolution equations,

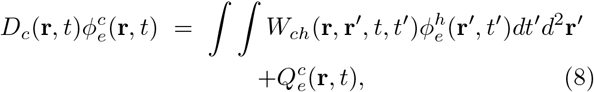

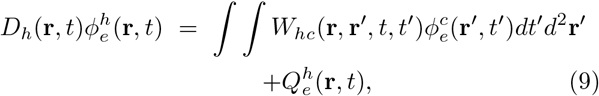

where the coupling functions *W*_*ch*_ and *W*_*hc*_ denote the hippocampus-to-cortex and cortex-to-hippocampus operators, respectively, constructed from the shared quasi-conformal coordinate system described above. The coordinates **r** and **r**′ denote target and source locations, respectively. Assuming that cortico-hippocampal transmission occurs on a macroscopic timescale comparable to corticothalamic transmission, we approximate the distributed axonal delays by a single characteristic delay (*τ*_*ch*_), which we set as *τ*_*ch*_ = *τ*_*c*_ = 0.04 s. This assumption allows for the temporal component of the interactions to be represented by Dirac delta function, *δ*(*t* − *t*′ − *τ*_*ch*_). Thus, the full spatiotemporal coupling function can be expressed as the product of spatial (*w*_*ch*_ and *w*_*hc*_) and temporal coupling terms, scaled by the coupling strengths *C*_*ch*_ and *C*_*hc*_,

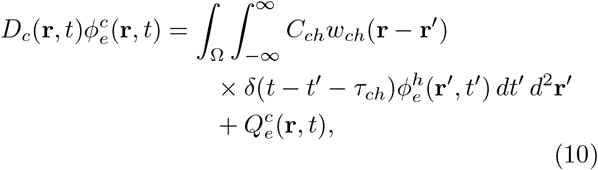

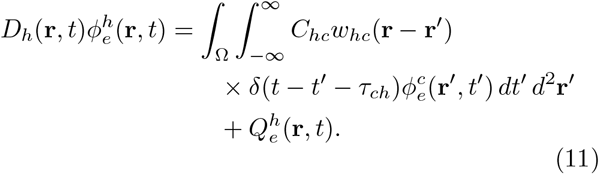

Smoothly decaying localised interactions are incorporated with a radially symmetric, exponentially decaying coupling kernel of the form 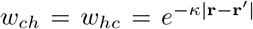, where *κ* governs the decay rate. By applying the identity, 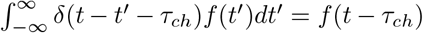, the time integral in the coupling equation simplifies to,

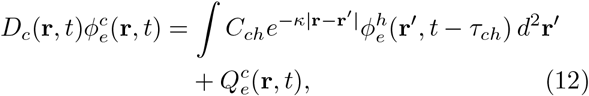

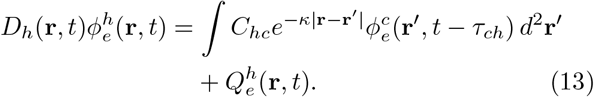

This integrated cortico-hippocampal system (Fig. 3a), allows insight into the impact of slowly increasing the inter-surface coupling strength (*C*_*ch*_ and *C*_*hc*_) on the unified cortico-hippocampal dynamics. Theoretical analyses suggest that bidirectional cortico-hippocampal coupling enhances cross-surface mode–mode interactions and leads to systematic shifts in the dominant system frequencies (Supplementary Materials S2). In agreement with these predictions, numerical simulations show that increasing coupling between the two surfaces initially sharpens and shifts the alpha and theta spectral peaks toward higher frequencies (Fig. 4a,b). These are characteristics of systems nearing critical transitions. Notably, the shifted and sharpened spectral peak of hippocampal theta (Fig. 4b, iii) aligns it more closely with empirical recordings (Fig. 2f and Fig. 4c).

**FIG. 4.**
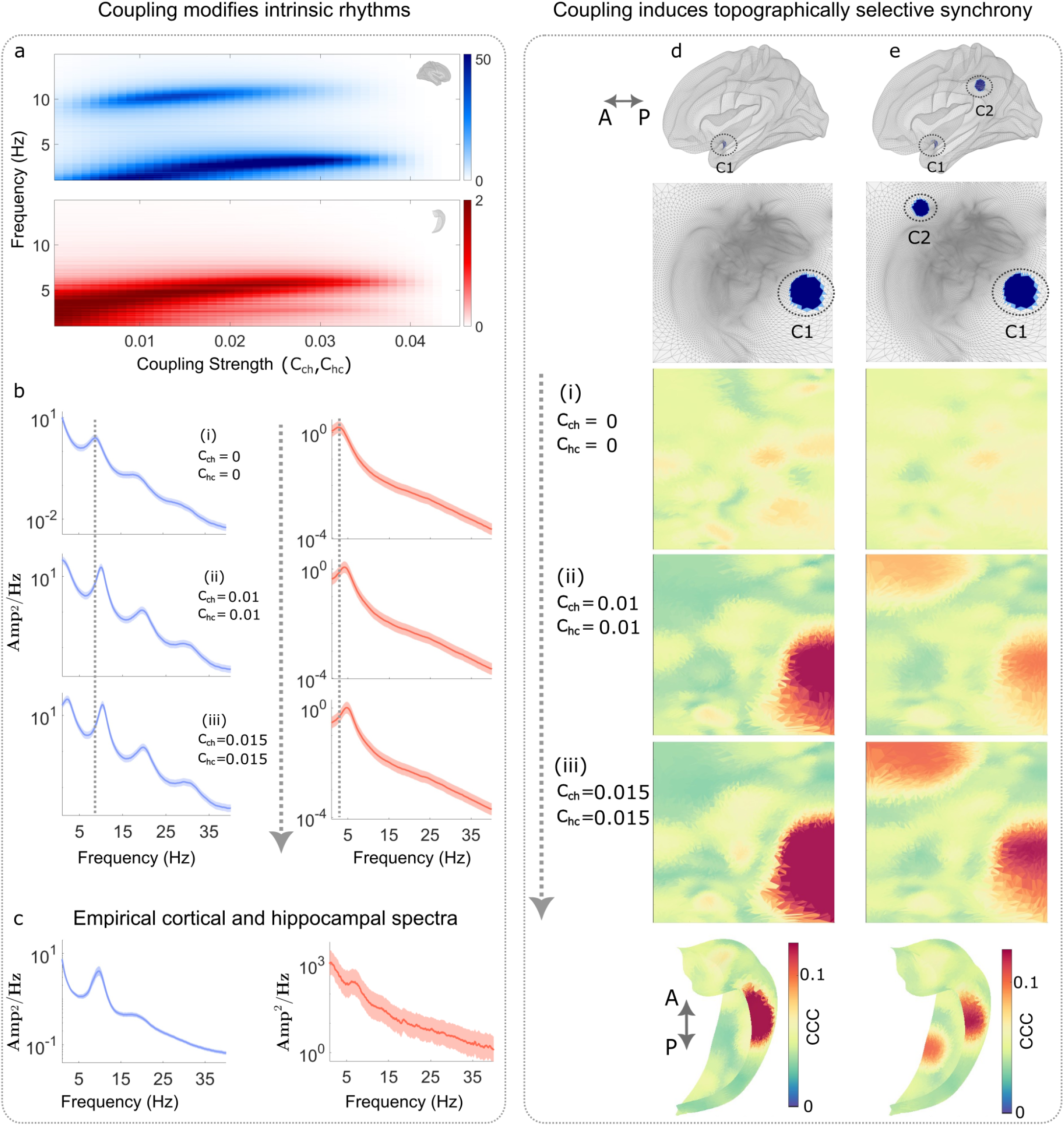
Influence of cortico-hippocampal coupling on spectral dynamics and spatial correlations. **a**, PSD heatmaps illustrating the shift of peak oscillation frequencies in the cortex (top, blue heatmap) and hippocampus (bottom, red heatmap) as a function of bidirectional coupling strength. **b**, Power spectral density (PSD) of cortical (blue) and hippocampal (red) eigenmode time series at three coupling strengths (*C*_*ch*_ = *C*_*hc*_ = 0 (**b -(i)**), 0.01 (**b -(ii)**), 0.015 (**b -(iii)**)), showing progressive sharpening and increase in dominant frequencies (arrows). **c**, Empirical spectra for the cortical alpha and hippocampal theta rhythms. **d**, Spatial specificity of cortico-hippocampal interactions, quantified by cross-correlation coefficients (CCC) between time series from a representative cortical region (blue patch) and all hippocampal vertices. In the uncoupled state (*C*_*ch*_ = *C*_*hc*_ = 0), correlations are weak and spatially diffuse. Stronger coupling (ii (*C*_*ch*_ = *C*_*hc*_ = 0.01), and iii(*C*_*ch*_ = *C*_*hc*_ = 0.015)) induces locally organized synchrony between topographically aligned cortical and hippocampal regions. **e**, Similar patterns of topographically organized cross-structure correlations emerge when analyses are extended to two cortical regions (namely, C1 and C2).

To further explore these trends, the strength of cortico-hippocampal coupling was increased in steps of 0.001 while keeping all other parameters for the corticothalamic and hippocampo-septal systems fixed within their respective alpha and theta rhythm regimes (Tables I, II). The spectral content of the resulting dynamics shows that stronger cortico-hippocampal coupling enhances synchrony and shifts activity toward faster frequencies on both surfaces (Fig. 4a). However, for excessive coupling between the two surfaces (*C*_*ch*_, *C*_*hc*_ *>*≈ 0.03), these peaks broaden and then disperse.

We next studied whether the structural mapping between the cortex and hippocampus yields a corresponding topographically specific functional mapping. Specifically, we checked whether the simulated cortical activity in the coupled cortico-hippocampal system aligns preferentially with topographically linked hippocampal regions and vice versa. To quantify the spatial specificity of cortico-hippocampal coupling, maximum lagged cross-correlations were calculated between the time series from hippocampal and cortical surfaces (see Methods). In the absence of cortico-hippocampal coupling, correlations between a single representative cortical region and hippocampal vertices remain low and spatially unstructured (Fig. 4d (i) and 4e (i)). With increasing corticohippocampal coupling (*C*_*ch*_ = *C*_*hc*_ = 0.01 and *C*_*ch*_ = *C*_*hc*_ = 0.015), this cortical region exhibits localized synchronization with specific hippocampal areas (Fig. 4d (ii) and 4d (iii); 4e (ii) and 4e (iii)). Notably, cortical and hippocampal regions occupying similar locations within their respective mapped domains display enhanced correlations. This pattern also reflects the anterior-posterior organization embedded in our coupling scheme. Extending this analysis to two representative cortical regions reveals that each synchronizes with a distinct, localized region of the hippocampal surface (Fig. 4e). These results confirm that cortico-hippocampal coupling promotes topographically organized synchrony while preserving local neighborhood structure, rather than inducing a global increase in coherence.

Taken together, the upward frequency drift, peak sharpening, and increased inter-system coherence are consistent with the approach to a critical transition. These changes, observed with increased coupling strength, suggest a potential mechanism by which the cortico-hippocampal system may shift from healthy dynamics into pathological states. In the following section, this hypothesis is tested by systematically varying the parameters that control excitability within the model. The resulting dynamics are then compared to empirical iEEG data acquired from human patients during epileptic seizures.

### E. Cortico-Hippocampal Seizures

Recurrent loops in large-scale neural architectures, such as the cortico-hippocampal system, are known to enhance mutual gain through positive feedback, thereby increasing the system’s susceptibility to dynamical instabilities [44, 71]. These instabilities have been associated with transitions to seizure-like states in both computational models and empirical studies [72, 73]. At the cellular level, glutamatergic neurons play a pivotal role in initiating and propagating epileptic seizures [74]. In line with this, glutamatergic transmission in the hippocampus has been implicated in driving hyperexcitability, with elevated extracellular glutamate levels observed in epileptogenic regions [75, 76]. During seizure-like transitions, increased functional coupling and phase synchrony between cortical and hippocampal networks have also been observed [77]. In our NFT model, the parameter 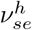 modulates the level of excitation in the hippocamposeptal feedback loop. The strength of excitatory cortico-hippocampal coupling is parameterised by *C*_*ch*_ and *C*_*hc*_. Together, these parameters define the excitatory feedback landscape across both local and long-range hippocampal loops and can be dynamically varied to model a representative hippocampal seizure consistent with focal onset Mesial Temporal Lobe Epilepsy (MTLE).

An almost ubiquitous feature of seizures is their dynamically changing spectral content [78, 79]. Individual hippocampal seizures exhibit substantial heterogeneity, but a recurring feature is the presence of spectral chirps, in which the dominant ictal frequency changes progressively over the course of the seizure [80]. To determine whether such dynamics are expressed across species and can be captured by the cortico–hippocampal model, we first compared seizure-related activity in human MTLE recordings, pilocarpine rat MTLE recordings [79], and PTZ-induced rat epilepsy recordings [81].

These empirical seizures share several prominent features: A dominant low-frequency ictal component with higher harmonics evident, plus increased synchronization within and between hippocampal and cortical systems (Fig. 5). Onset and termination are clinically annotated for the human MTLE seizures (Fig. 5a,b). The hippocampal and cortical spectrograms reveal seizure-related power concentrated in the theta range with a clear slowing of the peak frequency and its harmonics. The dynamic synchrony measure shows that the seizure is accompanied by increased hippocampus–hippocampus (H-H, red) and cortico–hippocampal (C-H, black) synchrony, whereas cortico–cortical (C-C, blue) synchrony increases only weakly and transiently (Fig. 5c). Similar patterns are present in the pilocarpine rat MTLE seizure (Fig. 5d–h). Multiple hippocampal and entorhinal subregions are recruited during the seizure, with hippocampal synchrony increased during the rhythmic theta component of the seizure activity. In the PTZ rat epilepsy dataset, group-averaged spectrograms also showed enhanced low-frequency power during the epileptic condition in both CA1 and M1 (Fig. 5j–n). This theta frequency activity during the seizure is also accompanied by increased cortico–hippocampal synchrony (Fig. 5n). To summarize, despite differences in species, recording modality, and seizure induction, these empirical seizures share convergent low-frequency ictal dynamics, including a progressive dominant-frequency trajectory and increased within- and cross-structure synchrony.

**FIG. 5.**
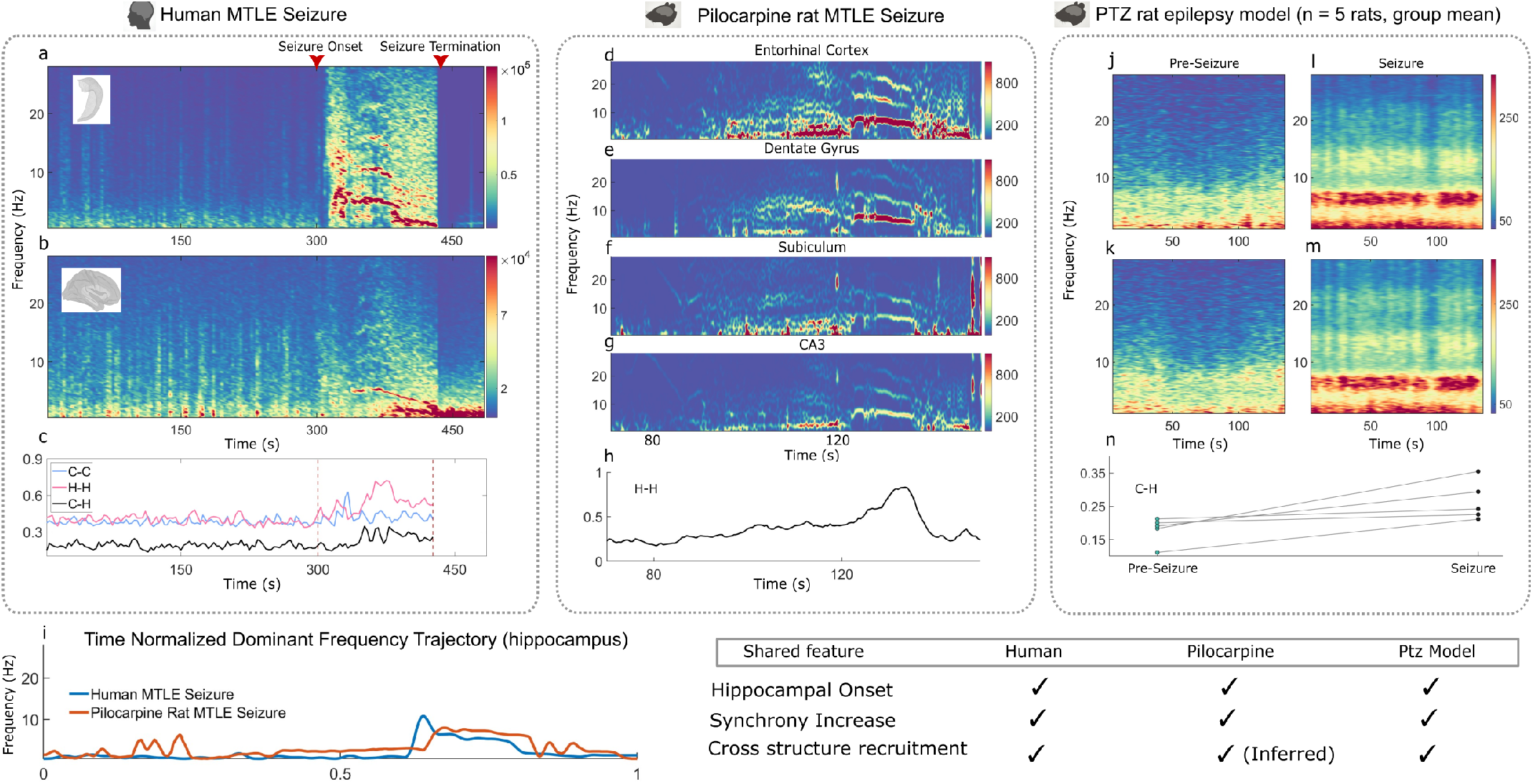
Mesial Temporal Lobe Epilepsy (MTLE) Seizures. **a** - **b** Representative intracranial EEG spectrograms from hippocampal (**a**) and cortical (**b**) recordings during a human MTLE seizure. Red triangles mark clinically annotated seizure onset and termination. (**c**) Dynamic synchrony measures computed between hippocampal–hippocampal (pink), cortico–cortical (blue), and cortico–hippocampal (black) electrode pairs. **d**–**h** Spectrograms from multiple entorhinal/hippocampal subregions (entorhinal cortex (**d**), dentate gyrus (**e**), subiculum (**f** ), and CA3 (**g**)) plus the corresponding hippocampus–hippocampus synchrony trajectory (**h**). (**j**–**m**) Group-averaged spectrograms from the PTZ rat epilepsy dataset across five rats. Spectrograms were computed from hippocampal CA1 (**j, l**) and motor cortex M1 (**k, m**) during healthy/pre-PTZ baseline (**j, k**) and PTZ-induced epileptic conditions (**l, m**). **n**, Cortico–hippocampal synchrony pre-seizure (left) and during seizure (right). **i**, Normalized trajectories of the dominant spectrogram frequency for the human and the pilocarpine rat MTLE seizures.

Next, we simulated hippocampal hyper-excitability, mimicking the corticohippocampal (human) and intra-hippocampal synchrony measures by increasing gain in the hippocampo-septal feedback loop, 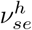 and increasing the bidirectional cortico–hippocampal couplings *C*_*ch*_ and *C*_*hc*_ (Fig. 6a–c). Hippocampal field activity, simulated in this way, exhibits a high amplitude oscillation, plus its harmonics, that closely mirrors the empirical human and pilocarpine rat MTLE seizures (Fig. 6d–f). Notably, the dominant hippocampal frequency in the simulated NFT closely matches the dynamics of the empirical seizures (Fig. 6g).

**FIG. 6.**
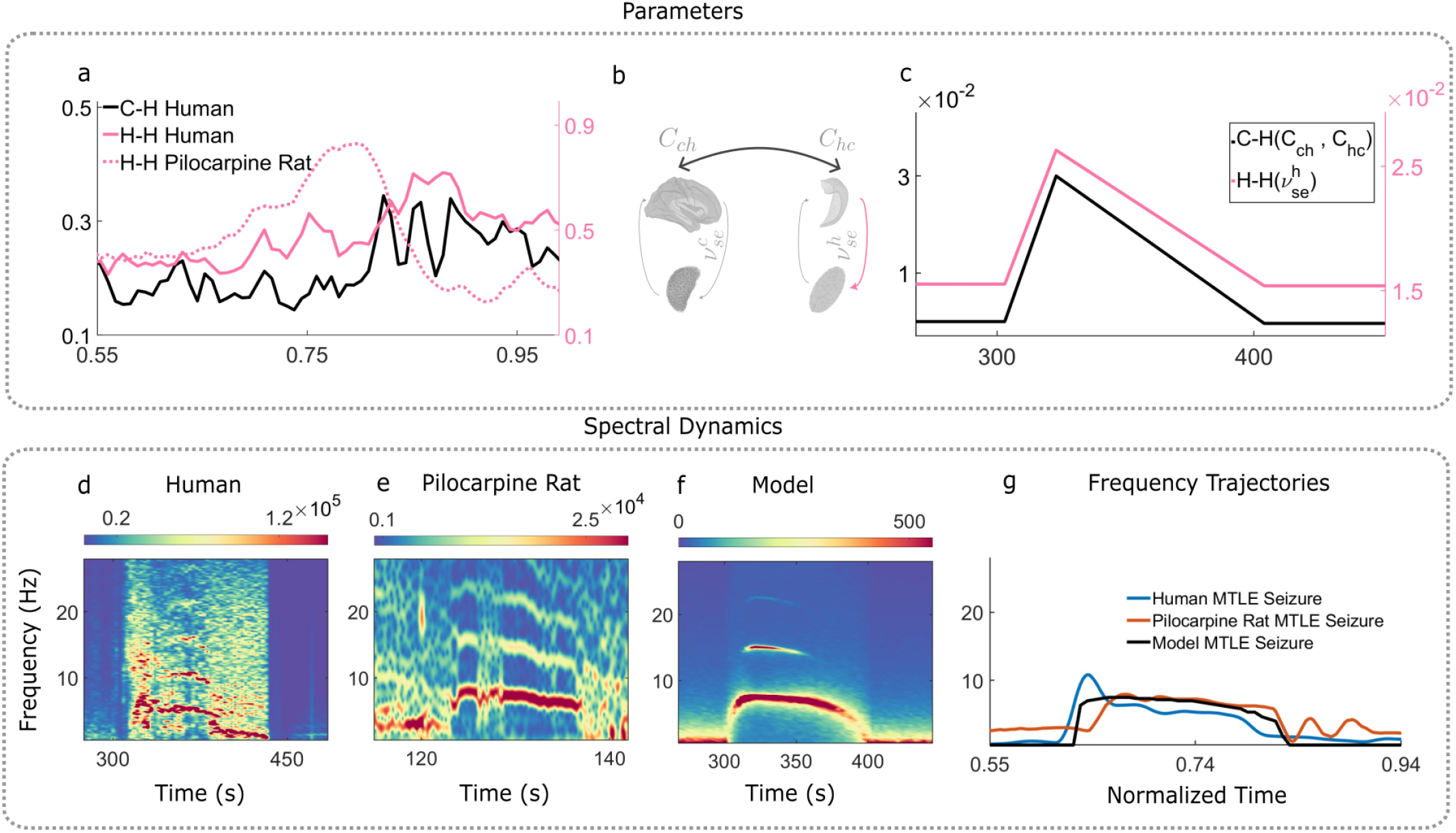
Convergent hippocampal frequency trajectories across human, rat, and model MTLE seizures. **a**, Dynamic synchrony measures from a representative human and a pilocarpine rat MTLE seizure, showing the evolution of cortico–hippocampal and hippocampus–hippocampus synchrony during the seizure epoch. **b**, Schematic summary of the coupled cortico–hippocampal model. **c**, Model parameter evolution trajectories for the bidirectional cortico–hippocampal coupling (*C*_*ch*_, *C*_*hc*_) and the excitatory gain parameter 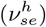 in the hippocampo-septal system. **d** - **f**, Zoomed spectrograms from human (**d**) and pilocarpine rat (**e**) hippocampal recordings, and the cortico–hippocampal model (**f** ), highlight a shared low-frequency ictal component near ∼ 7 Hz. The empirical examples correspond to seizure epochs described previously, while the model spectrogram is generated by increasing hippocampal excitability and cortico–hippocampal coupling. **g**, Time-normalized dominant hippocampal frequency trajectories for the human, pilocarpine rat and the model MTLE seizure.

A closer examination of the simulated NFT seizure reveals the spectral transition at seizure onset reorganizes multiscale activity across the hippocampal and cortical surfaces (Fig. 7). Prior to seizure onset, hippocampal activity is relatively low in amplitude with weak mode-mode coupling between hippocampal eigenmodes (Fig. 7a,b). During the seizure, as hippocampal power increases sharply and converges to the dominant theta band, mode-mode coupling strengthens broadly across hippocampal eigenmodes. This suggests that the increase in oscillatory power at seizure transition is accompanied by a spatial reorganization of hippocampal activity into more coherent cross-scale dynamics. Cortical field activity is also recruited during the simulated hippocampal seizure, although the cortical responses are weaker and more spatially selective. The cortical field shows a seizure-related increase in low-frequency power, with cortical mode–mode coupling increasing primarily among low-order modes (Fig. 7c,d). These low-order cortical modes represent broad spatial patterns, suggesting that hippocampal seizure activity can recruit large-scale cortical dynamics through cortico–hippocampal feedback. Cross-structure mode coupling supports this interpretation. During the simulated seizure epochs (Fig. 7), hippocampal modes became more strongly correlated with cortical modes, with the strongest coupling involving low-order cortical eigenmodes (Fig. 7e).

**FIG. 7.**
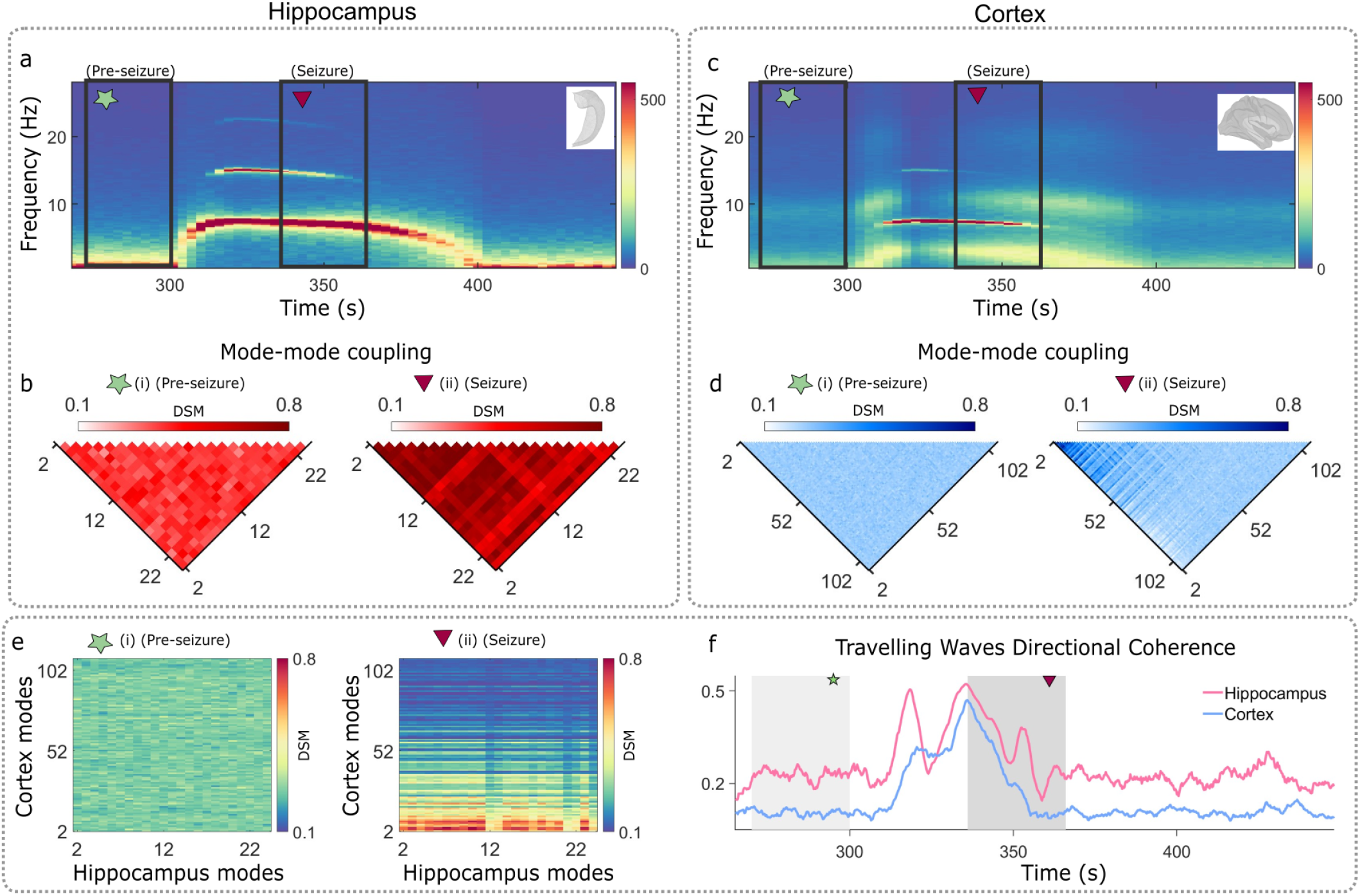
Simulated hippocampal-onset seizure. **a, c**, Average spectrograms of simulated hippocampal (**a**) and cortical (**c**) activity during the transition from the pre-seizure state to seizure activity. Green stars indicate the pre-seizure analysis window, and magenta triangles indicate the seizure window used for subsequent analyses. **b, d** Dynamic Synchrony Measure (DSM), computed as windowed maximum-lagged correlation between eigenmode time series. **e**, Cross-structure cortico–hippocampal mode coupling increases during the seizure, with hippocampal modes becoming more strongly correlated with cortical modes. **f**, Travelling-wave directional coherence (TWDC), computed as the band-limited alignment of local travelling-wave directions across surface vertices.

Clinical seizures frequently propagate across cortex [82, 83] and hippocampus [84, 85] as travelling waves. The corticohippocampal NFT provides an ideal platform to study these *in silico*. To achieve this, travelling-wave directional coherence (TWDC) was used to quantify the alignment of propagation patterns across cortex and hippocampus. TWDC in both structures increases during the seizure window, indicating that the transition into seizure activity is associated with more coherent travelling-wave organization across each surface (Fig. 7f). Together, these results show that increased hippocampal excitability and cortico–hippocampal feedback are sufficient to generate multiple hallmark features of MTLE-like seizure dynamics including low-frequency ictal activity, spectral chirping, enhanced within-hippocampal synchrony, increased cortico–hippocampal coupling, and coherent travelling-wave activity. The corticohippocampal NFT therefore provides a mechanistic bridge between empirical seizure signatures and the underlying dynamical reconfiguration of coupled hippocampal and cortical surface activity.

## III. DISCUSSION

Interacting fields evolving on distinct manifolds are encountered across many areas of physics. Their coupling typically relies upon a pre-existing geometrical relation between the manifolds on which they evolve. The fields may share a common coordinate system [86], be coupled across a common boundary [87], or interact through a mediating physical medium [88]. Large-scale brain dynamics introduce a different class of this problem. Continuous neural activity evolves on nonidentical anatomical manifolds that lack a natural point-to-point correspondence. Yet, their dynamics remain coupled through topographically organized projections that approximately preserve local neighborhood relationships [27, 89]. In this work, we address this problem by developing a coupled neural-field framework that enables spatially structured interactions between nonidentical manifolds. The resulting dynamics are shaped jointly by both the inter-manifold coupling and the feedback architecture of each manifold. At moderate coupling, the characteristic cortical alpha and hippocampal theta rhythms remain distinct. Their spectral peaks sharpen and shift, while coherence emerges selectively between topographically linked regions. On the other hand, when stronger coupling is combined with increased hippocampal excitability, the system enters a coherent low-frequency ictal regime. This transition is accompanied by enhanced cross-structure synchrony and more coherent travelling-wave activity across both surfaces.

This progression of dynamics distinguishes physiological coordination from pathological collective recruitment. In the physiological regime, the cortex and hippocampus remain dynamically differentiated. The characteristic cortical and hippocampal frequency spectra persist even as spatially selective synchronization emerges between the two manifolds. A moderately coupled cortico-hippocampal system therefore coordinates two distinct oscillatory fields through spatially selective cross-manifold synchronization. Such selective cortico-hippocampal coordination is consistent with empirical studies showing transient and regionally specific synchronization between hippocampal and cortical systems during cognition and memory processing [9, 10, 70]. In the pathological regime, increased hippocampal excitability is a well-established feature of MTLE. Seizure transitions are also accompanied by enhanced synchronization and the spatial propagation of neural activity [75–77, 83, 90]. In our model, increasing hippocampal excitability together with cortico-hippocampal coupling similarly recruits the two fields toward a shared low-frequency oscillation. This transition is also accompanied by stronger cross-structure synchronisation and increased directional coherence of travelling-wave activity across both manifolds.

These results, therefore, clarify the role of intermanifold coupling in the approach towards collective instability. Increasing cortico-hippocampal coupling sharpens and shifts the manifold-specific spectral peaks and increases region-specific coherence. The seizure state is not observed as a continuation of this frequency shift. A stronger coupling has to be accompanied by increased hippocampal excitability for the coupled feedback system to enters a distinct low-frequency oscillatory regime. Thus, inter-manifold coupling controls the approach to collective instability, while the dynamical state into which the system lands depends jointly on the coupling and local hippocampal excitability.

Our results separate the mechanisms governing the temporal and spatial organisation of cortico-hippocampal dynamics. The endogenous alpha and theta rhythms are set primarily by the feedback timescales, gains, and delays of the corticothalamic and hippocampo-septal systems. The low-frequency ictal rhythm emerges when these feedback architectures are driven into a strongly coupled, high-excitability regime. The neighbourhood-preserving coupling between these surfaces controls how this activity is exchanged across the spatial modes of the two systems. The coupling map therefore constrains where interactions can preferentially occur, while the field dynamics determine the strength, spectral content and mode structure of the resulting synchronization. Organised travelling activity is a recurrent feature of ictal propagation in both the cortex and the hippocampus [82, 83]. The increase in directional organisation of travelling-wave activity, without imposing a preferred propagation direction, links local hyperexcitability to the spatial organisation of seizure activity across both fields. Our framework therefore links oscillatory instability to its spatial expression across coupled anatomical mani-folds. More broadly, these results show how coupling between nonidentical active manifolds can reorganize collective dynamics in multilayer systems whose components differ in shape, scale, and intrinsic timescale.

An equally important contribution of the present work is methodological. Coupling spatially extended dynamics across distinct manifolds is challenging when the surfaces differ in geometry, scale, and discretisation. Such surfaces may also lack a natural point-to-point correspondence. We address this problem by mapping each surface independently to an aligned planar domain using quasi-conformal parameterisations. A distance-dependent kernel then transfers activity bidirectionally between the two fields. Quasi-conformal maps preserve topology and limit local angular distortion [58]. Hence, they can approximately retain local neighbourhood structure, without requiring identical meshes or vertexwise matching. We use these maps as a dynamical interface between continuous fields on nonidentical manifolds. Our construction, therefore, provides a controlled way to encode and compare hypotheses about how spatially separate systems exchange activity by testing alternative alignments, coupling kernels, transmission delays, and empirical connectivity constraints. Beyond the cortico-hippocampal system, the same strategy could be applied to other multilayer systems with continuous dynamics on non-matching curved surfaces.

Contextualizing our results within the field of large-scale brain modeling highlights the distinct explanatory potential of neural-field approaches [31]. High-fidelity reconstructions such as Spaun [91] and the Digital Brain [92] offer biological detail but are computationally expensive and challenging to invert. Data-driven foundation networks [93] occupy the other end of the spectrum, achieving strong predictive accuracy but offering limited insight into underlying mechanisms. The present framework occupies a complementary middle ground, combining computational efficiency with the ability to derive closed-form solutions that treat the brain as a physical dynamical system. This, in turn, links parameters such as gain and inter-manifold coupling to specific neurobiological processes, enabling principled interpretation of macroscopic brain activity and linking geometry and emergent dynamics in a theoretically transparent manner.

Our framework also makes several testable predictions. It predicts that ictal recruitment should lead to an increase in low-frequency power along with a coordinated rise in cross-structure mode coupling. This rise is reflected in an increased directional wave coherence within both cortical and hippocampal surfaces. The model further predicts a dissociation between temporal and spatial effects of coupling. Spatially randomising the coupling operator may preserve a collective oscillatory instability while disrupting topographically selective synchronisation and organised mode recruitment. These predictions can be tested using simultaneous cortical and hippocampal recordings and subject-specific anatomical reconstructions.

The present work also brings a specific focus on cortico-hippocampal mean field dynamics, foregrounding the important role of the hippocampus in adaptive cognition and pathological seizure states. We initiated hippocampal seizures via an increase in the coupling and gain parameters of the hippocampus. The increase in gain enhances excitation in the system and simulates the gluta-matergic release observed during seizure onset [74]. Future extensions could incorporate seizure-specific meanfield models [72, 94, 95] or explicit metabolic mechanisms of seizure generation [96]. These additions would provide a more direct link between local pathophysiological processes and large-scale brain dynamics.

A complementary set of extensions could incorporate the anatomical specificity and individualisation of the framework. First, the classical neural field theory assumes an isotropic and uniform synaptic kernel approximated by the exponential distance rule (EDR) for the axonal projections. Incorporating anatomically informed long-range connections would enable insight into the influence of second-order attributes of the connectome on large-scale brain dynamics. Second, integrating subject-specific cortical and hippocampal surfaces reconstructed from magnetic resonance imaging (MRI), together with individualized connectomes, could transform the framework into a personalized digital twin [95]. Finally, the hippocampal theta in our framework is generated through hippocampo-septal feedback, consistent with the established role of the medial septum in theta generation [56, 97]. However, hippocampal dynamics are also shaped by interactions with structures including the entorhinal cortex and thalamus [24–26, 98]. Incorporating these pathways would enable the framework to examine how multiple afferent systems jointly regulate physiological rhythms and pathological recruitment.

## IV. METHODS

### Neural Field Theory (NFT) parameters

The parameters for the corticothalamic system were adopted from previously published papers where the corresponding NFT predicted and reproduced key features of cortical dynamics [43, 44, 48, 59] (Table I).

Thalamocortical projections are relatively sparse and topographically organized [99, 100]. In contrast, the medial septum forms dense and widespread connections with the hippocampus [66, 101, 102]. To reflect these anatomical differences, the synaptic gain parameters in the hippocampal-septal loop were empirically scaled relative to those of the corticothalamic system. Furthermore, because the hippocampus and septum are in close anatomical proximity, with fast-acting GABAergic and cholinergic projections mediating hippocampal theta rhythms [56, 57], a shorter delay (*τ*_*h*_ = 5 ms) was employed than the corresponding corticothalamic delay (*τ*_*c*_ = 40 ms). This parameter set captures the stronger local coupling and reduced conduction delays characteristic of the hippocampo-septal circuitry while preserving the mathematical structure of the cortico-thalamic NFT (Table II).

### A. Eigenmode decomposition and Integration

To integrate the neural field equations on the cortical and hippocampal surfaces, the spatial dependence of each field was represented using the geometric eigenmodes of the corresponding anatomical surface. For each structure *a* ∈ {*c, h*}, the geometric eigenmodes 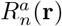 are defined as the eigenfunctions of the Laplace–Beltrami operator,

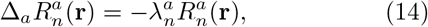

where Δ_*a*_ denotes the Laplace–Beltrami operator on the cortical (*a* = *c*) or hippocampal (*a* = *h*) surface, and 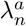 is the eigenvalue associated with the *n*th eigenmode. These eigenmodes are determined by the geometry of the corresponding surface and form an orthogonal basis for representing smooth scalar fields defined on that surface.

The propagating excitatory field can therefore be expanded as a weighted sum of geometric eigenmodes,

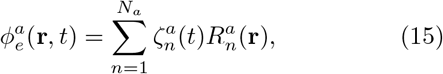

where 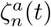 denotes the time-dependent amplitude of the *n*th eigenmode and *N*_*a*_ is the number of modes retained for structure *a*. The field 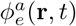 evolving on the cortical or the hippocampal surfaces is sustained by the net excitatory and inhibitory input these surfaces receive from local cortical or hippocampal populations and the thalamic or the septal nuclei. These inputs are embedded in the source term 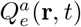. The source term can also be represented in the same geometric eigenmode basis as follows,

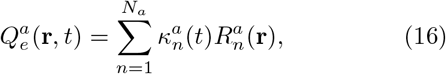

where 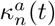 is the projection of the source activity onto the *n*th geometric eigenmode. Substituting Eqs. (15) and (16) into the neural field equation and using Eq. (14) gives

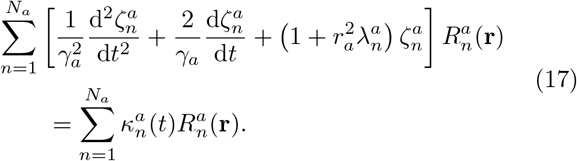

Projecting this expression onto each eigenmode and using their orthogonality yields a system of temporal equations,

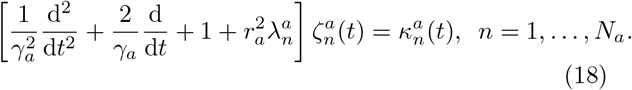

This reduces the original spatially extended neural field PDE to a finite-dimensional ODE system for the modal amplitudes. The linear spatial propagation operator is diagonal in the geometric eigenmode basis. However, the modal dynamics are not generally independent because the coupling between modes is retained through the source coefficients 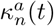, which depend on the non-linear population responses, delayed feedback pathways, and inter-manifold coupling terms. Consequently, the complete model forms a coupled system of nonlinear delay differential equations in the modal amplitudes and the associated population variables.

To efficiently capture the dominant spatial patterns while maintaining computational tractability, the field 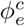 on the cortical surface was represented using the first 110 geometric eigenmodes. The highest retained cortical mode corresponds to a minimum represented spatial wavelength of approximately 40 mm, consistent with recent empirical findings [5]. The hippocampal field was represented using the first 25 eigenmodes, corresponding to a minimum represented spatial scale of approximately 3 mm. The different truncation levels account for the markedly different physical dimensions of the cortical and hippocampal surfaces and define the spatial band-width retained within each field representation. Following previous work [43, 73], the characteristic range for thalamic nuclei and the medial septum was set to zero, thereby suppressing local propagation across these brain areas. This simplification allows thalamic populations to be modeled as discrete nodes rather than continuous fields, in contrast to the surface-based treatment of the cortex and the hippocampus. This eigenmode-based formulation enables region-specific control over spatial autocorrelation by adjusting the number of modes used in each structure.

### B. Coupling between the cortex and the hippocampus

The cortical and hippocampal surfaces used in this study are population-averaged anatomical manifolds. To simulate dynamic cortico-hippocampal interactions, a bidirectional coupling scheme was introduced, treating both the cortex and the hippocampus as continuous two-dimensional neural sheets. Let **r** denote spatial locations on the surface of interest (either cortex or hippocampus), and **r**′ denote locations on the surface where the field originated. This convention enables a consistent formulation of interactions across the coupled cortico-hippocampal system.

As per Results Section D, the coupled neural field dynamics are governed by,

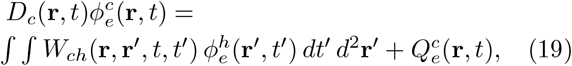

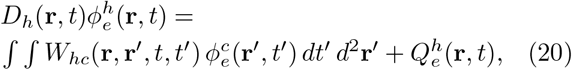

where *D*_*c*_ and *D*_*h*_ are the wave operators defined for the cortex and hippocampus, respectively. The functions *W*_*ch*_ and *W*_*hc*_ describe the spatiotemporal coupling from hippocampus to cortex and vice versa. Source terms 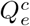 and 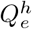 denote the excitatory firing rate and act as an input to the cortical/hippocampal NFT equations.

To retain spatial specificity and analytical tractability, we assume the coupling is uniform and isotropic and thus only depends on spatial and temporal offsets,

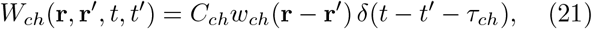

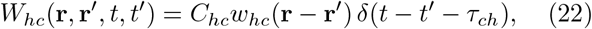

where *w*_*ch*_(·) and *w*_*hc*_(·) are spatial kernels, and *τ*_*ch*_ is the fixed transmission delay. In all simulations, the cortico-hippocampal transmission delay was set equal to the corticothalamic delay listed in Table I, *τ*_*ch*_ = *τ*_*c*_ = 0.04 s.

To construct these spatial coupling kernels, we independently map the cortical and hippocampal surfaces to two-dimensional domains using quasi-conformal mapping [58]. A quasi-conformal map is a homeomorphism from a 2D manifold (surface) embedded in R^3^ to a planar domain, preserving local topology and introducing only bounded angle distortion (with Beltrami coefficient *µ*(*z*) satisfying |*µ*(*z*)| *<* 1). For each surface *S*_*a*_ (*a* ∈ {*c, h*} for cortex and hippocampus), we obtain a bijective map

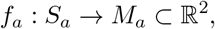

ensuring a one-to-one correspondence between the curved surface in 3D and its 2D flattened, planar representation. Notably, this procedure does *not* define a bijection between cortical and hippocampal surfaces them-selves, as their geometries and discretizations generally differ. Instead, the flattening simply provides a consistent 2D domain for subsequent kernel construction and enables proximity-based coupling, where interactions between surfaces are determined by distances between vertices in their respective 2-D embeddings.

The coupling strength between a cortical vertex *c*_*i*_ and a hippocampal vertex *h*_*j*_ is then defined using an exponentially decaying function of Euclidean distance,

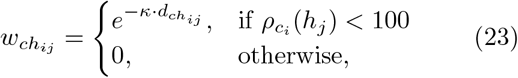

where 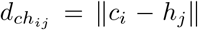, *κ* is the decay constant, and 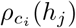 is the rank of hippocampal vertex *h*_*j*_ by distance from cortical vertex *c*_*i*_. Each cortical vertex receives weighted input from its 99 nearest hippocampal neighbors in this shared Euclidean space, and vice versa. Under the assumption of approximately uniform intervertex spacing, this enforces a consistent projection area across the surfaces. Similarly sized cortical patches influence each hippocampal location, and vice versa. This implementation supports spatially selective field transport between surfaces.

This coupled cortico-hippocampal system was integrated using the same eigenmode-based approach described above. The cortical and hippocampal surfaces were each decomposed into geometric eigenmodes, and their respective wave equations were solved in this space. The coupling therefore acts as an additional inter-manifold source term, and this combined input (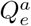 and the coupling input) was projected into the eigenmode basis. The coupled wave equations were also solved in the eigenmode space. Thalamic and septal populations remained point-based and were integrated independently at each location, consistent with the assumption of zero spatial range. This scheme enabled consistent and computationally efficient simulation of the coupled surface–point system.

### Power spectral density estimation

The spectral content of cortical and hippocampal activity was quantified using a one-sided power spectral density (PSD). For a sampled time series *x*[*n*] with sampling frequency *F*_*s*_, the signal was first detrended and multiplied by a Hann window *w*[*n*]. The discrete Fourier transform of the windowed signal was

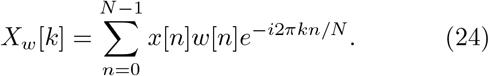

The two-sided periodogram was normalized by the sampling frequency and the energy of the applied window,

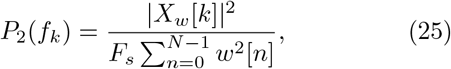

where the discrete frequency bins are

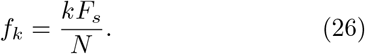

For real-valued signals, the one-sided PSD was obtained by retaining frequencies between 0 and *F*_*s*_*/*2 and multiplying the positive-frequency components by two, excluding the Nyquist component when present.

### Spatial Specificity of Cortico-Hippocampal Coupling

To assess the spatial specificity of cortico-hippocampal coupling, cross-correlation coefficients (CCCs) were calculated between time series from cortical regions of interest (ROIs) and individual hippocampal vertices. Because a single cortical vertex carries negligible functional relevance, a small cortical region—comprising a central vertex and its 99 nearest neighbors was used to approximate a localized population (Fig. 4d and 4e, blue patches). The maximum lagged cross-correlation between each hippocampal vertex and all 100 vertices in the selected cortical region was computed and averaged to yield a single correlation score per hippocampal vertex. Repeating this analysis for every hippocampal vertex produces a spatial map of how strongly each hippocampal location synchronizes with the chosen cortical patch. Representative cortical ROI (Fig. 4d and 4e), was defined as the 100 nearest cortical vertices to a selected seed point. That is, for a hippocampal vertex *h*_*k*_ and a cortical vertex *v*_*i*_, the lagged cross-correlation 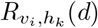 was calculated over the entire time series, within a lag range of ±1 s,

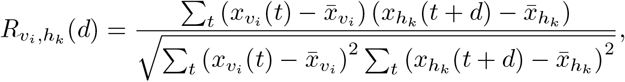

where 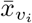 and 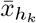 are mean values of the respective time series and *d* is the temporal lag.

The maximal absolute cross-correlation was then calculated as,

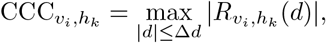

where Δ*d* = 1 s.

For each hippocampal vertex *h*_*k*_, the average CCC across all the vertices in the cortical ROI vertices was calculated as,

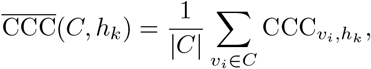

where |*C*| = 100 is the number of cortical vertices in the ROI *C*. This produced hippocampal surface maps of 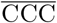, reflecting the strength of coupling between each cortical ROI and the entire hippocampus. Since the results demonstrated in Fig. 4 are for a constant cortico-hippocampal coupling strength, system dynamics were stationary, allowing the use of these static measures without temporal windowing.

### Dynamic Synchrony Measure

Time-varying synchrony was quantified using a Dynamic Synchrony Measure (DSM) based on windowed maximum-lagged cross-correlation. The same core definition was applied to empirical electrophysiological recordings and simulated eigenmode time series.

Signals were band-pass filtered between 1 and 15 Hz. DSM was calculated with 50% overlap between windows. Within each window, the mean was removed independently from each signal before calculation of the cross-correlation. For two signals *x*_*i*_(*t*) and *x*_*j*_(*t*) within temporal window *w*, the demeaned signals are

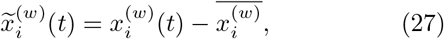

and

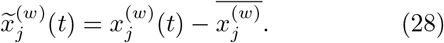

The normalized lagged cross-correlation was then calculated as

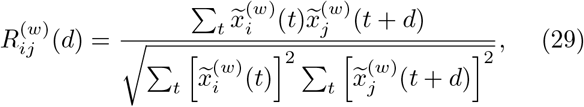

where *d* denotes the temporal lag.

The DSM for a given pair of signals and temporal window was defined as the maximum absolute normalized cross-correlation within a lag range of ±1 s,

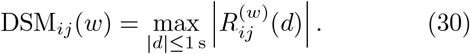

Thus, DSM quantifies the magnitude of the strongest lagged linear relationship between two band-limited signals within each temporal window, irrespective of whether the corresponding correlation is positive or negative.

For empirical within-structure analyses, DSM was calculated for all included electrode pairs within the hippocampus (H–H) or cortex (C–C). Pairwise DSM values were averaged within each temporal window to obtain a structure-level synchronization trajectory,

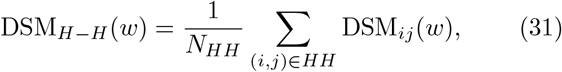

with an analogous definition for DSM_*C*−*C*_(*w*). Cross-structure cortico-hippocampal synchronization was calculated across included cortical–hippocampal electrode pairs,

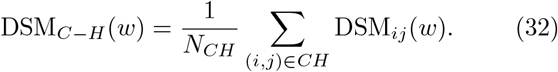

For simulated data, DSM was calculated between geometric eigenmode coefficient time series. For the mode– mode synchronization matrices, DSM was calculated independently in each temporal window and subsequently averaged across windows for every pair of modes. The initial quarter of the simulated time series was discarded before stationary mode–mode analyses to remove initial transient dynamics.

No Hilbert-phase transformation was used in the DSM calculation. This ensures that the empirical and simulated analyses share the same cross-correlation-based definition and avoids applying a linear correlation measure directly to wrapped phase angles.

### Spectral Content Analysis via Short-Time Fourier Transform

To visualize the temporal evolution of spectral content during seizures, time–frequency spectrograms were calculated using the short-time Fourier transform (STFT). Individual time series were analyzed using a Hann window for both the human and the rodent datasets. Since the sampling frequency varied across datasets, the corresponding window and FFT lengths in samples were determined independently from the native sampling frequency of each dataset. The native sampling frequencies were 500 Hz for the human MTLE recordings, 2000 Hz for the pilocarpine rat recordings, and 1000 Hz for the simulated cortical and hippocampal time series.

Spectrograms were calculated separately for individual empirical electrodes or simulated time series and were subsequently averaged within each anatomical region to obtain representative cortical or hippocampal spectrograms.

### C. Dominant-frequency trajectory extraction and temporal normalization

To characterize the evolution of the dominant low-frequency component during seizure activity, dominant-frequency trajectories were extracted from the hippocampal spectrograms. For the pilocarpine rat recording, electrophysiological signals were sampled at *F*_*s*_ = 2000 Hz. The sampling frequency for the human MTLE seizure was 500 Hz, and the simulation time series were sampled at 1000 Hz. Each signal was demeaned and band-pass filtered between 0.5 and 45 Hz using a fourth-order Butter-worth filter applied in the forward and reverse directions to avoid phase distortion.

Time-resolved spectral power was then calculated using a short-time Fourier transform. For the pilocarpine rat data, spectrograms for recordings from CA3, entorhinal cortex, dentate gyrus, and subiculum were averaged to get a representative spectrogram for the pilocarpine rat hippocampus.

The power at each frequency was first smoothed across three consecutive spectrogram time bins using a moving mean. The dominant-frequency trajectory was then defined as the frequency containing the maximum smoothed power within the range 0.5–15 Hz. The human and rat seizures lasted for different durations. To compare dominant-frequency trajectories across seizures with different absolute durations, different seizure trajectories were mapped onto a common normalized time coordinate. The normalized seizure trajectories were then linearly interpolated onto a shared grid of 450 equally spaced points covering the interval 0 ≤ *τ* ≤ 1. Only the temporal coordinates were normalized and the dominant-frequency values remained expressed in Hz. The resulting curves therefore describe dominant hippocampal frequency as a function of normalized displayed time, enabling comparison of the relative spectral evolution of human and pilocarpine rat MTLE seizures despite their different absolute durations.

### D. Travelling-wave directional coherence

To quantify the spatial organization of travelling-wave activity during the simulated seizure, we calculated travelling-wave directional coherence (TWDC) separately across the cortical and hippocampal surfaces. Vertex-level cortical and hippocampal activity was reconstructed from the corresponding geometric eigenmode coefficients. For each structure *a* ∈ {*c, h*}, the reconstructed field is given by,

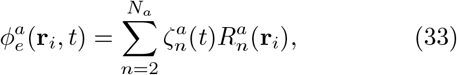

where 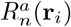 denotes the *n*th geometric eigenmode evaluated at vertex *i*, and 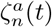 is its time-dependent coefficient. The first spatially constant eigenmode was excluded, and all remaining available eigenmodes were retained.

Local spatial neighbourhoods were defined independently on the cortical and hippocampal surfaces using the two-dimensional quasi-conformal coordinates described above. For each vertex, the 12 nearest vertices in the corresponding quasi-conformal domain were identified using Euclidean distance, thereby forming a local *k*-nearest-neighbour graph. The reconstructed vertex time series were band-pass filtered between 3 and 10 Hz, corresponding to the dominant low-frequency activity observed during the simulated seizure. The analytic signal was then obtained using the Hilbert transform,

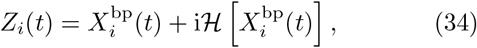

where 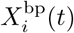 is the band-pass-filtered signal at vertex *i* and *H*[·] denotes the Hilbert transform. The instantaneous phase and amplitude were respectively defined as,

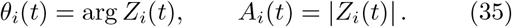

The local travelling-wave direction was estimated from the spatial phase gradient. For a vertex *i* with neighbouring vertices *j* ∈ *N*_*i*_, the coordinate displacement matrix was defined from,

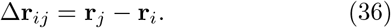

Because phase is a circular variable, phase differences were wrapped to the interval [−*π, π*] according to,

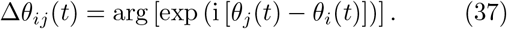

The local phase-gradient vector **k**_*i*_(*t*) was then obtained by solving the linear approximation,

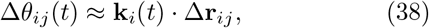

in a least-squares sense. In matrix form,

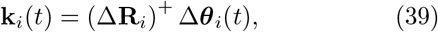

where Δ**R**_*i*_ contains the spatial displacement vectors between vertex *i* and its neighbours, Δ***θ***_*i*_(*t*) contains the corresponding wrapped phase differences, and (·)^+^ denotes the Moore–Penrose pseudoinverse. Vertices with fewer neighbours than the dimension of the coordinate space, or with zero-distance neighbours, were excluded from the local gradient calculation.

The phase-gradient vectors were normalized to obtain local unit direction vectors,

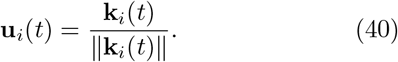

Vertices for which ∥**k**_*i*_(*t*)∥ *<* 10^−8^ were excluded because the local propagation direction was not reliably defined. The vector **u**_*i*_(*t*) points in the direction of increasing phase. The physical direction of wave propagation may equivalently be represented by −**u**_*i*_(*t*). However, this sign convention does not affect the directional-coherence magnitude used here.

TWDC was defined as the magnitude of the spatially averaged local direction vectors,

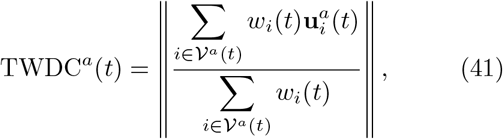

where *V*^*a*^(*t*) denotes the set of vertices with valid phase-gradient estimates at time *t*. In the present analysis, amplitude weighting was not used, such that *w*_*i*_(*t*) = 1 for every valid vertex. This ensures that increases in oscillatory amplitude during the seizure do not directly increase the estimated directional coherence. TWDC was not evaluated when fewer than 10 vertices contained valid local direction estimates.

TWDC values range between zero and one. Values approaching zero indicate that local phase-gradient vectors are weakly aligned or spatially heterogeneous across the surface, whereas values approaching one indicate that travelling-wave directions are coherently aligned across a common direction. Cortical and hippocampal TWDC were computed independently and therefore quantify within-surface travelling-wave organization rather than directional coherence directly between the two structures. TWDC was evaluated every 0.25 s. The cortical and hippocampal fields were reconstructed and analysed in consecutive 10-s temporal blocks. Each block included 5 s of temporal padding on either side during band-pass filtering and Hilbert-transform estimation, while TWDC values were retained only from the central, unpadded interval. The resulting TWDC trajectories were smoothed using a 5-s moving-average window. The pre-seizure and seizure intervals shown in Fig. 7 corresponded to 270– 300 s and 336–366 s, respectively.

## ACKNOWLEDGEMENTS

M.B. and R.P. were supported by the National Health and Medical Research Council (APP2008612). This study was supported by QIMR Berghofer Clinician Research Collaboration Grant Award 6626 and a Collaboration Agreement between QIMR Berghofer and Mater Research. M.B. acknowledges funding support of the Advanced Queensland Innovation Partnership (AQIP12316-17RD2). The authors acknowledge the past and present team members of Mater Epilepsy Unit and Mater Research (Brisbane) for technical and/or administrative support. J.C.P. acknowledges support from the Australian National Health and Medical Research Council (ID: 2034000) and Monash FMNHS Early Career Research Excellence Program. The authors acknowledge the facilities and scientific and technical assistance of the National Imaging Facility (NIF), a National Collaborative Research Infrastructure Strategy (NCRIS) capability, at the Hunter Medical Research Institute Imaging Centre, University of Newcastle. The funding bodies had no role in study design, analysis, decision to publish, or preparation of the manuscript.

## AUTHOR CONTRIBUTIONS

R.P. and M.B.: Study conception and development of the modelling framework. R.P.: Numerical simulation. P.A.R., R.P., and M.B.: Theoretical development and mathematical formalism. S.S.: Access to human epilepsy data, preprocessing (filtering, down-sampling, re-referencing), and extraction of de-identified data derivatives. S.S. and J.A.R.: Generation of figures showing epilepsy time traces. R.P. and M.B.: Provision of custom scripts for signal visualization. R.P.: Visualization of the spectral density and correlation time series and provision of scripts for visualization of all other numerical and empirical data. J.D.: Provision of the hippocampal surface meshes. R.P., M.B., J.A.R., J.C.P., S.S., S.D., A.B., N.K., J.D., J.M.S., P.A.R. and A.F. contributed to the interpretation of results and provided domain expertise. R.P. and M.B. drafted the initial version of the manuscript. All authors reviewed, edited, and approved the final version of the manuscript.

## DATA AND CODE AVAILABILITY

The empirical data analysed in this study were obtained from publicly available datasets, restricted clinical recordings, and data shared directly by the original investigators.

Resting-state scalp EEG data used to characterise cortical alpha activity in Fig. 1 were obtained from the publicly available OpenNeuro SRM resting-state EEG dataset [62]. Human hippocampal intracranial EEG data used as an example of hippocampal theta rhythm in Fig. 2 were obtained from the publicly available Open-Neuro TSS iEEG dataset [69].

The pilocarpine rat mesial temporal lobe epilepsy recording used in Fig. 5 and 6 was provided directly by Dr Maxime Lévesque upon request. This data corresponds to the recording underlying Figure 2A of Lévesque et. al. [79]. The recording was shared as a MATLAB file containing signals from CA3, entorhinal cortex, dentate gyrus, and subiculum, sampled at 2,000 Hz with a 500-Hz hardware low-pass filter. As permission to redistribute the underlying raw recording was not explicitly granted, these data should be requested from the original data provider.

The PTZ-induced rat epilepsy recordings used to make the spectrograms in Fig. 5 and 6 were obtained from the publicly available intracranial EEG dataset reported by Ghanaei and Erfanian [81]. These recordings contain hippocampal CA1 and motor-cortical M1 activity from five rats under baseline and PTZ-induced epileptic conditions.

De-identified human intracranial EEG seizure recordings used as an example of human MTLE seizure in Figs 5 and 6 are from the Mater Advanced Epilepsy Unit, Mater Hospital, Brisbane. These recordings were originally acquired for clinical purposes and cannot be made publicly available because of participant consent, privacy, and institutional ethics restrictions. Future access to these data will require local ethics approval and a data sharing agreement with the Research Governance Offices of QIMR Berghofer and Mater Research, Brisbane.

Custom MATLAB and EEGLAB scripts used for data analysis, as well as neural field simulation codes are available openly online [103].

