## Supplemental PDF for "Geometry-Constrained Coupling Drives Critical Dynamics Across Neural Manifolds"

### OVERVIEW

This supplementary material includes theoretical and numerical methodological details for the results presented in the main manuscript. Parameter choices, simulation protocols, and data processing methods are also provided.

### S1. ANALYSIS

#### S1.1. Cortical and Hippocampal Eigenmodes

The geometric eigenmodes of the cortical and hippocampal surfaces are computed by solving the Laplace–Beltrami eigenvalue problem. These eigenmodes provide a natural basis for modeling field dynamics on curved two-dimensional manifolds. Each eigenmode represents a spatially distinct pattern determined by the geometry of the surface [1, 2]. For the cortical surface, the fsLR 32k population-average template mesh from the Human Connectome Project (HCP) is used. The medial wall is removed to produce a continuous, simply connected domain. A pre-processed anatomical mesh defines the hippocampal surface [3, 4]. No masking or exclusion is required for this surface.

The cotangent-weighted Laplace–Beltrami operator $\mathbf{L}$ is constructed using finite-element discretization [5]. For each edge $(i, j)$, the weights are given by,

$$\mathbf{L}_{ij} = \frac{1}{2}\left(\cot\alpha_{ij} + \cot\beta_{ij}\right), \qquad \text{(S1)}$$

where $\alpha_{ij}$ and $\beta_{ij}$ are the angles opposite edge $(i, j)$ in its two adjacent triangles. At boundary edges, where only one adjacent triangle exists, the weight reduces to,

$$\mathbf{L}_{ij} = \frac{1}{2}\cot\alpha_{ij}. \qquad \text{(S2)}$$

This formulation provides a discrete approximation of the continuous Laplace–Beltrami operator and implicitly applies Neumann boundary conditions.

A Voronoi-area-based mass matrix $\mathbf{M}$ is computed to account for non-uniform vertex areas and to improve numerical stability. The generalized eigenvalue problem,

$$\mathbf{L}\psi_n = \lambda_n \mathbf{M}\psi_n \qquad \text{(S3)}$$

is solved using MATLAB's sparse eigensolver `eigs`. The first 110 eigenmodes are retained for the cortical surface, and the first 25 eigenmodes are retained for the hippocampal surface. This ensures an approximate match of the higher spatial frequency.

#### S1.2. Quasi-Conformal Mapping of Cortical and Hippocampal Surfaces

In order to establish a systematic coupling scheme between the cortex and hippocampus, a common coordinate system is needed in which both surfaces can be consistently compared. Since the cortex and hippocampus exist as distinct, non-overlapping manifolds in three-dimensional space, directly computing interactions between them is non-trivial due to differences in their intrinsic geometries, curvatures, and spatial embeddings. To overcome this, Quasi Conformal Mapping (QCM) is used. The cortex simplifies to a connected open surface after the medial wall is removed. The hippocampal surface is already a connected open surface. Both surfaces

*
†

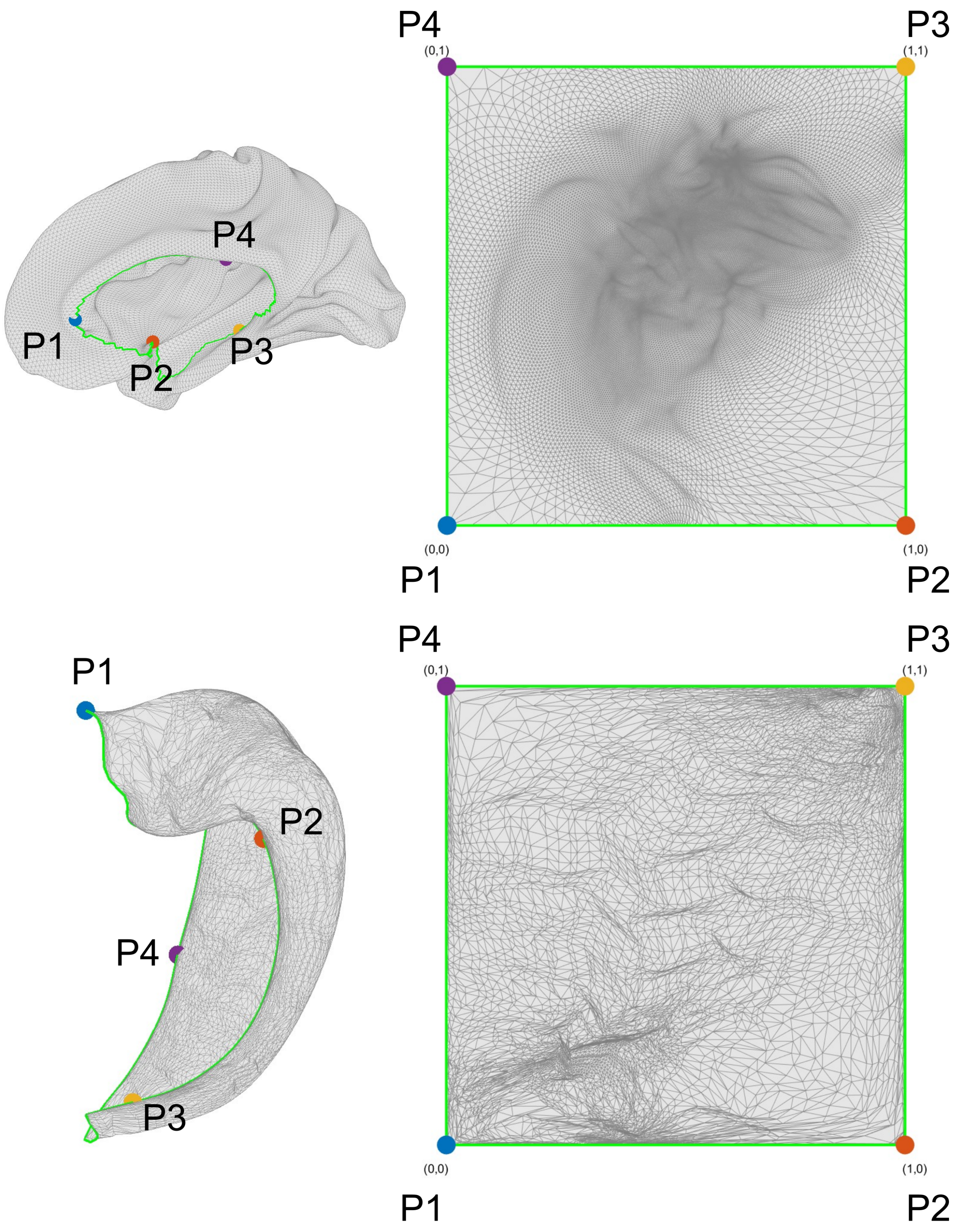


FIG. S1. Quasi-conformal mapping of cortical and hippocampal surfaces to a shared Euclidean domain (a unit square). The cortical and hippocampal 3-D surfaces (left panels) are flattened into a common 2-D square domain (right panels) using quasi-conformal mapping. Four anchor points (P1–P4), selected along the cortical medial wall and the hippocampal boundary, are mapped to the square's corners to preserve anterior–posterior and medial–lateral orientation. The medial wall boundary of the cortex and the hippocampal boundary are highlighted in green to show continuity in the flattened domain.

can therefore be flattened to a planar domain via QCM (Fig. S1). The algorithm by Meng et al. [6] is used to perform the mapping by solving the Beltrami equation,

$$\frac{\partial f}{\partial \bar{z}} = \mu(z)\frac{\partial f}{\partial z} \tag{S4}$$

where $f : \mathbb{C} \to \mathbb{C}$ represents the mapping from the 3-D surface to a 2-D domain, and $\mu(z)$ is the complex-valued Beltrami coefficient. The coefficient $\mu(z)$ describes the amount and direction of local distortion, while the constraint $|\mu(z)| < 1$ ensures that the mapping remains quasi-conformal and orientation-preserving. The triangulation of the original surface meshes is also preserved during mapping. This maintains local vertex-face relationships and enables accurate reconstruction of surface geometry after flattening. A landmark-based boundary condition is enforced by mapping four anchor points to the corners of a square (Fig. S1), thereby aligning corresponding cortical and hippocampal boundaries in the 2-D domain. Two of the anchor points are placed towards the anterior and posterior ends of both the cortical and hippocampal surfaces, preserving the anterior–posterior gradient in the mapping and ensuring biologically consistent cortico-hippocampal alignment (Fig. S1).

QCM approximately preserves neighborhood relationships between points with local distortions in scale. The key advantage of QCM lies in its ability to represent cortical and hippocampal activity fields within the same coordinate space, making it possible to define a proximity-based interaction kernel without dealing with the complexities of their original 3-D embeddings. By aligning the two surfaces in a common domain, a spatially coherent and mathematically tractable coupling scheme between the two surfaces can be implemented (Fig. S2). This implementation defines the interaction strength between points on the cortical and hippocampal maps using an adaptive nearest-neighbor kernel. An adaptive kernel is necessary because the local distortions caused by QCM introduce non-uniform vertex densities, causing some regions to be compressed more than others. Using fixed Euclidean cutoff distances for a spatial kernel would make the kernel highly uneven in the original 3-D embedded cortical and hippocampal structures. Hence, a coupling range is instead defined by selecting the nearest 99 neighbors of the source vertex closest to the target vertex in the Euclidean space. Such a coupling ensures that when mapped back to the original 3-D geometry, the interaction range remains approximately uniform, assuming an approximately constant inter-vertex spacing.

To implement this coupling, the field coming to a target vertex $r$ (Fig. S2 d, h, l, p) is computed by aggregating contributions from a region $r'$ on the source surface. This region $r'$ is defined as the 99 nearest neighbors to the mapped target point in the 2-D domain (Fig. S2 b, f, j, n). As illustrated in Fig. S2 a, b, e, and f, the coupling kernel compensates for local compression or expansion introduced by the mapping, ensuring that the effective projection area remains approximately constant in the original 3-D geometry (Fig. S2 a and e). Fig. S2 a - h show hippocampus-to-cortex field transport, while Fig. S2 i - p depict cortex to hippocampus field transport. Together, these panels summarize how the QCM framework enables smooth and invertible coupling between the two surfaces, preserving local topology and supporting spatially localized inter-regional interactions.

S3 a–h demonstrate the effect of this coupling on field transport. Panels a–d show a smoothly varying field on the hippocampal surface being projected to the cortex. The transported field retains spatial coherence, reflecting the smoothing and the topology and topography preserving property of the coupling scheme. Fig. S3 e–h show the transport of a more spatially complex field from hippocampus to cortex with fine-scale details being smoothed while preserving the overall structure of the field.

### S1.3. Individualized seizure dynamics for additional patients

#### *1. Patient Information*

The present study analyzed de-identified iEEG data obtained from a previously established cohort of patients with drug-resistant epilepsy who underwent stereotactic depth electrode implantation at the Mater Advanced Epilepsy Unit, Mater Hospital, Brisbane, as part of their clinical evaluation. Electrode placement was determined solely on clinical grounds. For the current analyses, we selected seizure recordings from patients with electrode contacts in both the cortical regions and the hippocampus (patients P1, P2 and P3). The original data acquisition was approved by the Human Research Ethics Committees of the Mater Hospital and the QIMR Berghofer Medical Research Institute, and all patients provided informed consent at the time of collection.

For details regarding access to these data, preprocessing, extraction of de-identified data derivatives and generation of figures, please refer to the Author Contribution and Data and Code Availability sections in the main manuscript.

#### *2. Data acquisition and preprocessing*

The iEEG data analyzed here were originally acquired during stereo-EEG monitoring at the Mater Advanced Epilepsy Unit, Mater Hospital, Brisbane. Intracranial depth electrodes (DIXI Medical; 10–15 contacts, 1 mm diameter, 1.5 mm spacing) were stereotactically implanted, with electrode implantation scheme guided by the epileptologist. Signals were recorded on a Neurofax EEG-1200 system (Nihon Kohden, Japan) at a sampling rate of 1 kHz. These data were preprocessed offline using

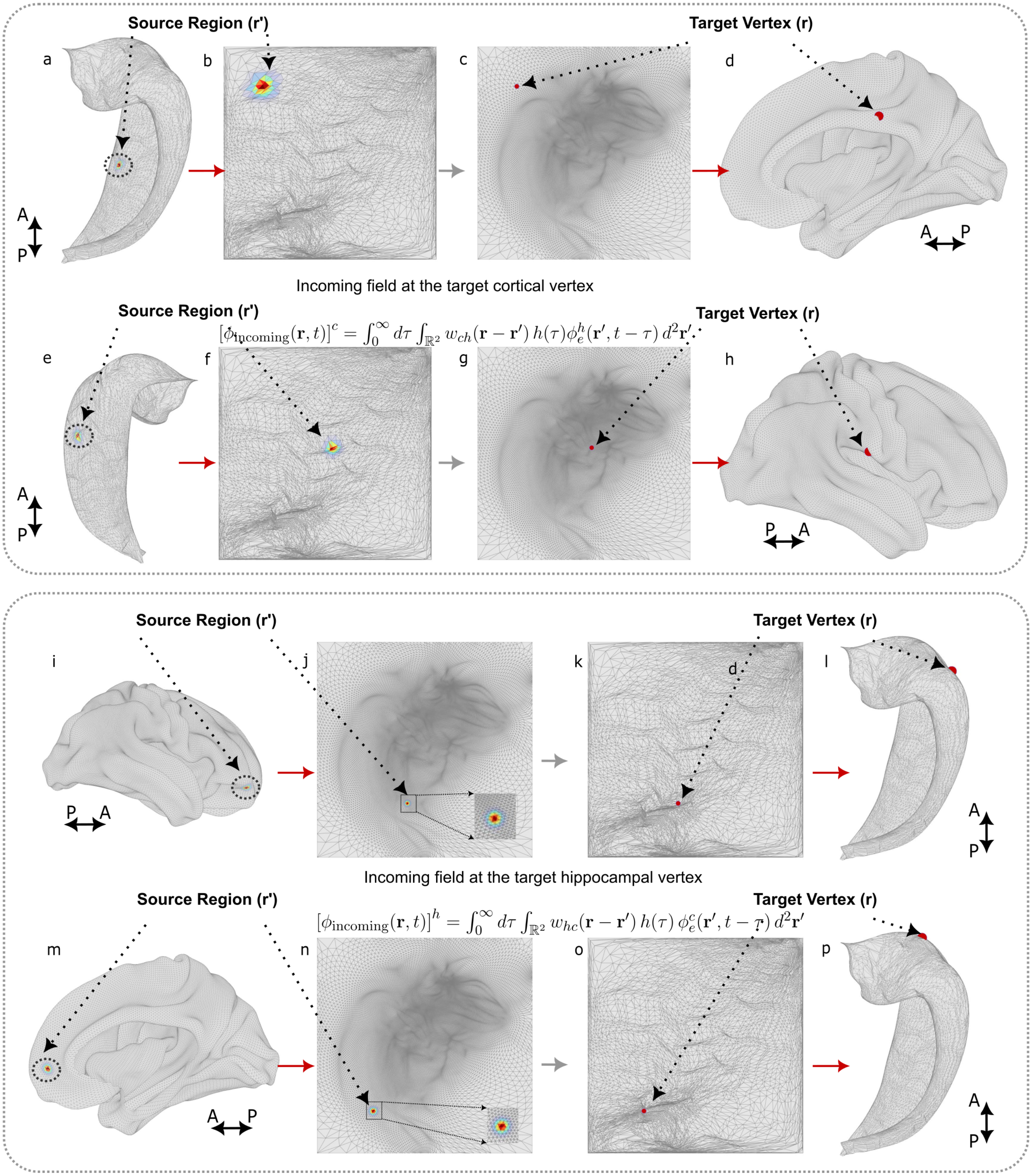


FIG. S2. Schematic of cortico-hippocampal coupling between the source and the target. Activity fields are mapped between cortical and hippocampal surfaces using a proximity-preserving, distance-dependent coupling kernel. (a–h) illustrate hippocampus-to-cortex field transport: each cortical vertex $(r)$ receives a weighted field as input from a localized hippocampal region $(r')$ defined by an exponentially decaying function of Euclidean distance $e^{-\kappa\|r-r'\|}$. (i–p) show cortex-to-hippocampus field coupling with an analogous mapping. The incoming field at each target vertex is computed as $\phi_{\text{incoming}}(r,t) = \int_{\Omega_2} C_{hc} w_{hc}(r-r')\, \phi_e^c(r', t-\tau_{ch})\, d^2r'$, where $w_{hc}(r-r')$ represents the spatial kernel and $\tau_{ch}$ accounts for finite conduction delays. Source region and target vertices are shown on both the original 3-D surfaces and their 2-D quasi-conformal mappings.

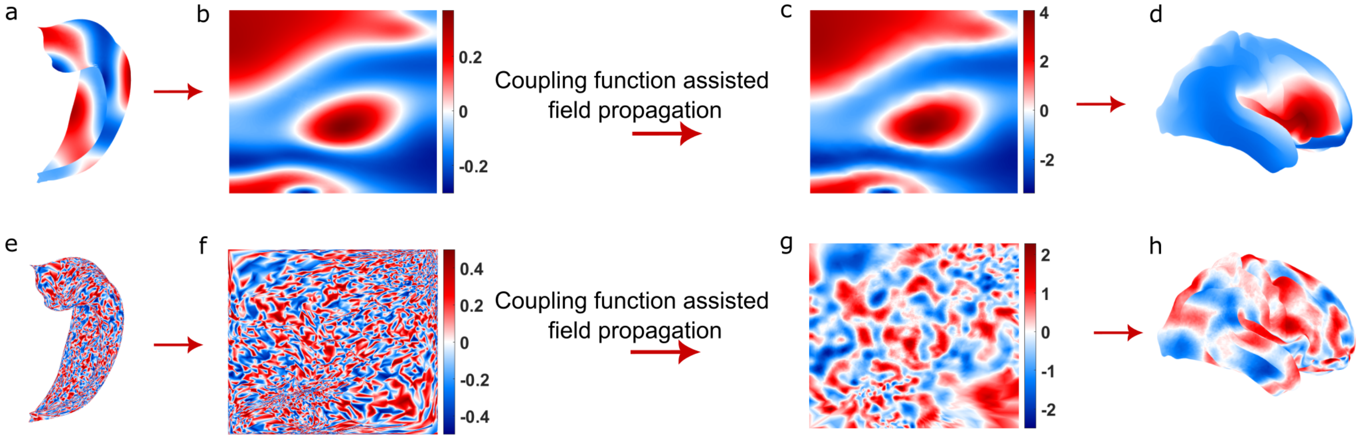


FIG. S3. Cortico-hippocampal field transport and its effects on spatial coherence. a–d Propagation of a smooth hippocampal field to the cortical surface, demonstrating preservation of spatial coherence during mapping. (e–h) Transport of a complex, random hippocampal field to the cortex, where fine-scale details are smoothed while overall structure is retained. The coupling operation applies an exponentially decaying spatial kernel to aggregate source-region contributions and includes a fixed time delay to account for finite propagation speed.

EEGLAB and custom MATLAB routines. Signals were downsampled to 500 Hz and band-pass filtered between 0.5 and 50 Hz with a zero-phase finite impulse response filter. All analyses here used a bipolar re-referencing scheme.

#### 3. *Simulating individual seizure dynamics*

To check for the model's ability to reproduce patient-specific seizure dynamics, we simulated seizure dynamics for three patients. For each case, time-resolved measures of intra- and inter-structural synchrony were used to guide parameter evolution in the model. These empirically observed synchrony trajectories were then used to guide parameter evolution in the model, enabling us to reproduce key spectral features of the seizures, most notably the characteristic spectral chirps shown in Fig. S4, Fig. S5 and S6.

The patients were adults aged between 30 and 50 years (2 females, one male) all with seizures originating in the medial temporal region (including hippocampus, parahippocampal gyrus, entorhinal cortex, and amygdala).

### S2. THEORY

For biologically motivated cortico-hippocampal coupling within NFT, it is important to incorporate the known topographic organization of projections between the two structures. Cortical–hippocampal connections are spatially and topographically organized. Anterior cortical regions preferentially couple to anterior hippocampal regions, and posterior regions to posterior hippocampal regions [7, 8]. Functional neighborhoods are preserved across regions, and long-range connections also follow spatial continuity. To capture these features, a shared coordinate frame is introduced using quasi-conformal mapping, which can provide a proximity-preserving correspondence between cortical and hippocampal surfaces. In this framework, the anatomical organization implies that local neighborhoods in one structure should map to local neighborhoods in the other, and smooth field transport emerges as a consequence of these constraints. In the following sections, a general framework is set up by considering the case of uniform coupling between the cortex and the hippocampus. However, an anisotropic and non-uniform cortico-hippocampal coupling can be treated within the same framework by modifying the spatial kernel that is used for the coupling scheme.

#### S2.1. Coupled Sheets

The activity fields $\phi_e^c$ and $\phi_e^h$ in two neural sheets obey equations of the form,

$$D_c(\mathbf{r},t)\phi_e^c(\mathbf{r},t) = \iint W_{ch}(\mathbf{r},\mathbf{r}',t,t')\phi_e^h(\mathbf{r}',t')\,dt'd^2\mathbf{r}' + Q_e^c(\mathbf{r},t), \tag{S5}$$

$$D_h(\mathbf{r},t)\phi_e^h(\mathbf{r},t) = \iint W_{hc}(\mathbf{r},\mathbf{r}',t,t')\phi_e^c(\mathbf{r}',t')\,dt'd^2\mathbf{r}' + Q_e^h(\mathbf{r},t). \tag{S6}$$

where $t$ and $t'$ denote time and $\mathbf{r}$ and $\mathbf{r}'$ denote spatial coordinates on different surfaces depending on the direction of coupling (Fig. S2). $\mathbf{r}$ is used for points on the

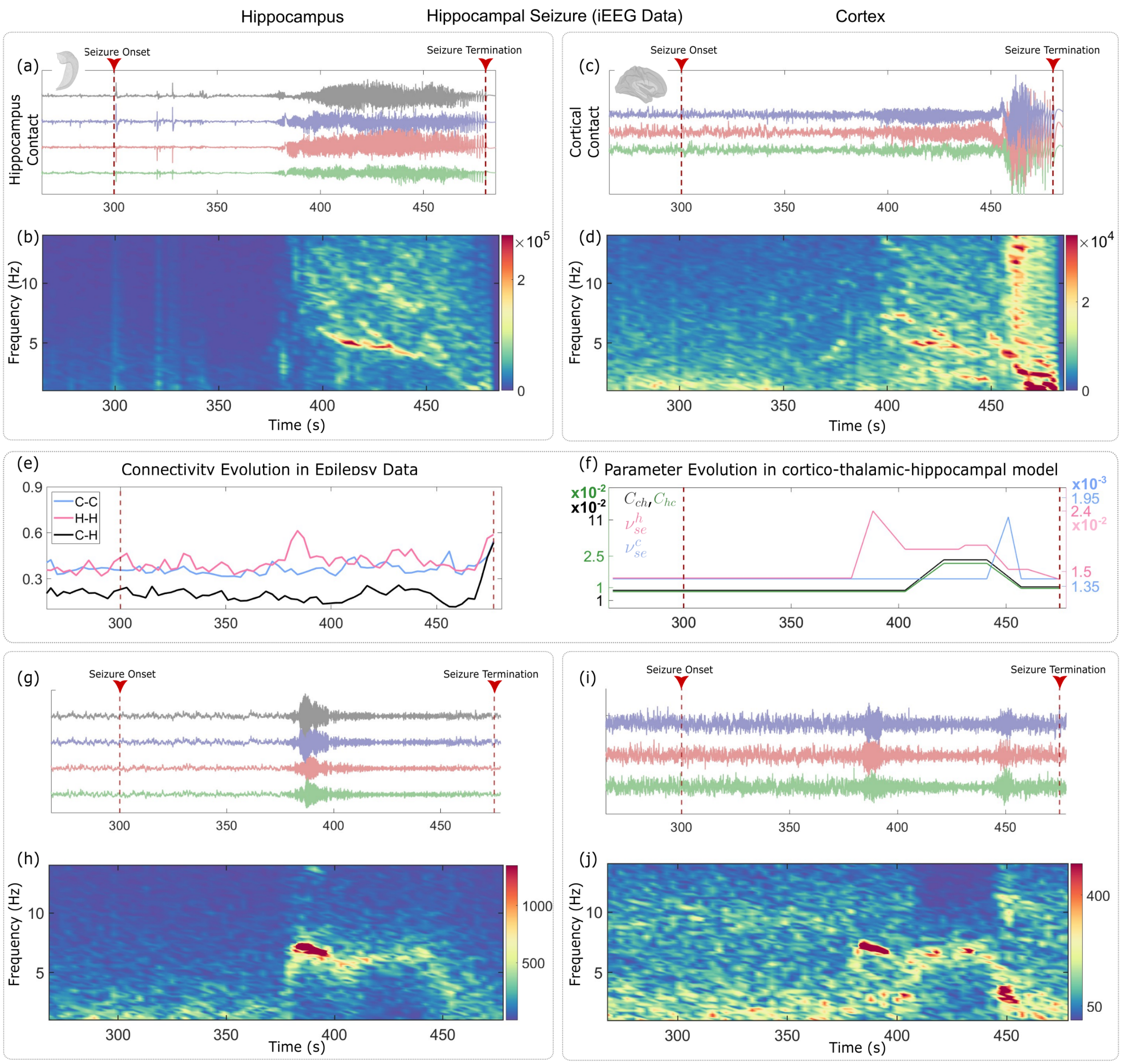


FIG. S4. Empirical and simulated seizure dynamics. (a) Preprocessed intracranial EEG (iEEG) traces from four hippocampal electrodes and (c) three cortical electrodes. Dotted red lines mark the onset and termination of the seizure as annotated by an epileptologist. (b,d) Average Short-time Fourier transform (STFT) spectrogram of the hippocampal electrodes (b) and the cortical electrodes (d). (e) Evolution of synchrony across unique hippocampal electrode pairs (pink), cortical electrode pairs (blue), and cortico–hippocampal electrode pairs (black). (f) System parameter trajectories inferred from the dynamic synchrony evolution in (e): hippocampal/cortical excitatory drive ($\nu_{se}^{h}/\nu_{se}^{c}$) and cortico-hippocampal coupling ($C_{ch}$ and $C_{hc}$). (g,i), Simulated time series from representative hippocampal (g) and cortical (i) vertices. (h, j), Simulated spectrogram from a representative hippocampal (h) and cortical vertex (j).

target surface (the surface onto which the field is being mapped) and $r'$ for points on the source surface (the surface at which the field originates). Integration is performed on the source variable $\mathbf{r}'$ to compute the values in the target variable $\mathbf{r}$. $D_c$ and $D_h$ are differential operators, and the coupling functions $W_{ch}$ and $W_{hc}$ can be quite general. $Q_e^c$ and $Q_e^h$ are source terms embodying the feedback activity coming from the reticular and relay nuclei of the thalamus and the cortical feedback activity.

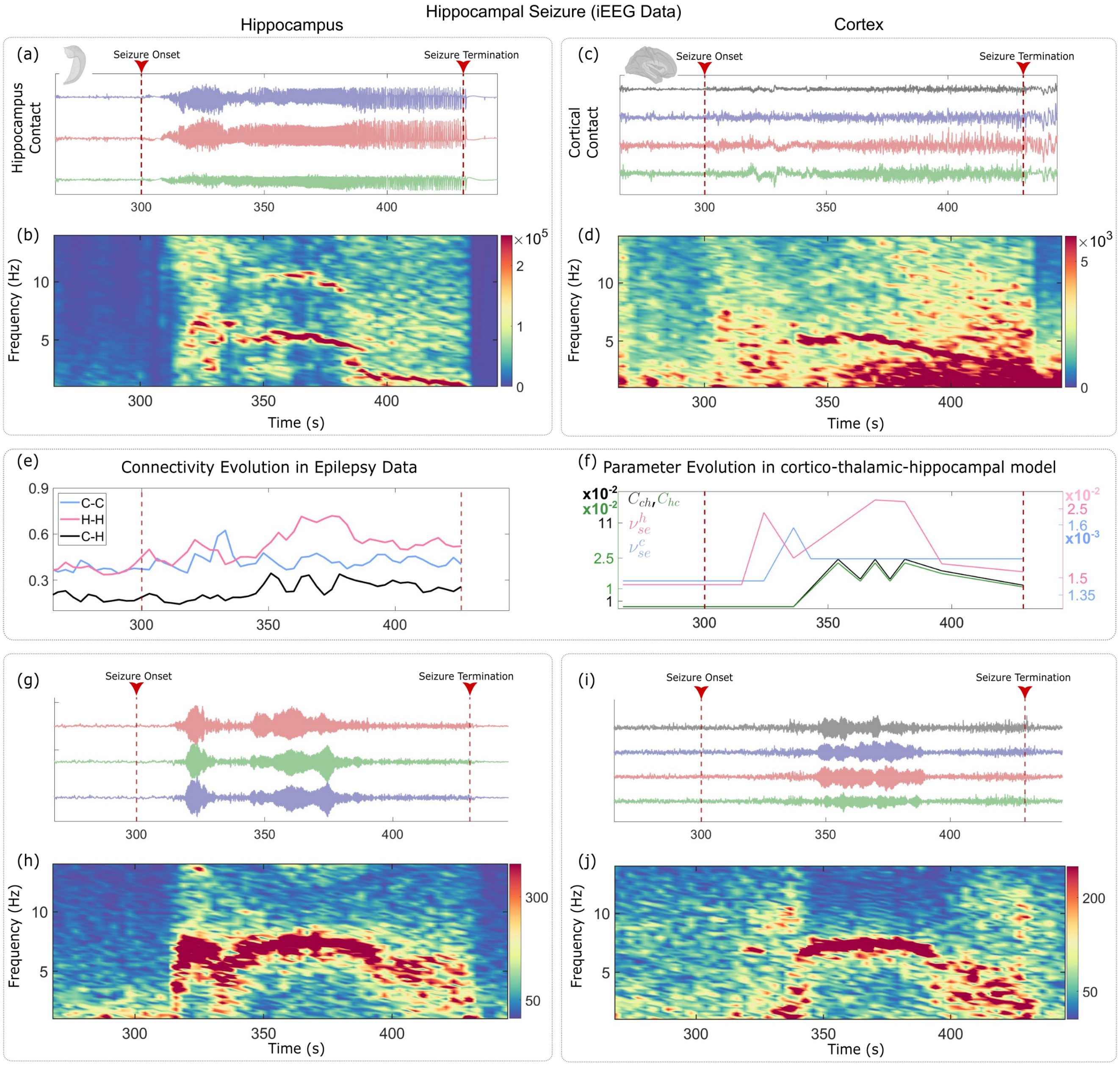


FIG. S5. Empirical and simulated seizure dynamics. (a) Preprocessed intracranial EEG (iEEG) traces from three hippocampal electrodes and (c) four cortical electrodes. Dotted red lines mark the onset and termination of the seizure as annotated by an epileptologist. (b,d) Average Short-time Fourier transform (STFT) spectrogram of the hippocampal electrodes (b) and the cortical electrodes (d). (e) Evolution of synchrony across unique hippocampal electrode pairs (pink), cortical electrode pairs (blue), and cortico–hippocampal electrode pairs (black). (f) System parameter trajectories inferred from the dynamic synchrony evolution in (e): hippocampal/cortical excitatory drive ($\nu^h_{se}/\nu^c_{se}$) and cortico-hippocampal coupling ($C_{ch}$ and $C_{hc}$). (g,i). Simulated time series from representative hippocampal (g) and cortical (i) vertices. (h, j), Simulated spectrogram from a representative hippocampal (h) and cortical vertex (j).

#### S2.2. Uniform Coupling

The coupling function has the same functional form for all the target vertices. This ensures that a theoretical analysis is possible, whether the mapping is isotropic or anisotropic in space. In this work, the coupling function only depends on differences in position and time (Fig. S2). Hence, in order to define coupling from the hippocampus to the cortex, an exponentially weighted field from a hippocampal region is mapped to a cortical vertex. Incorporating this in the governing wave equations S5 and S6,

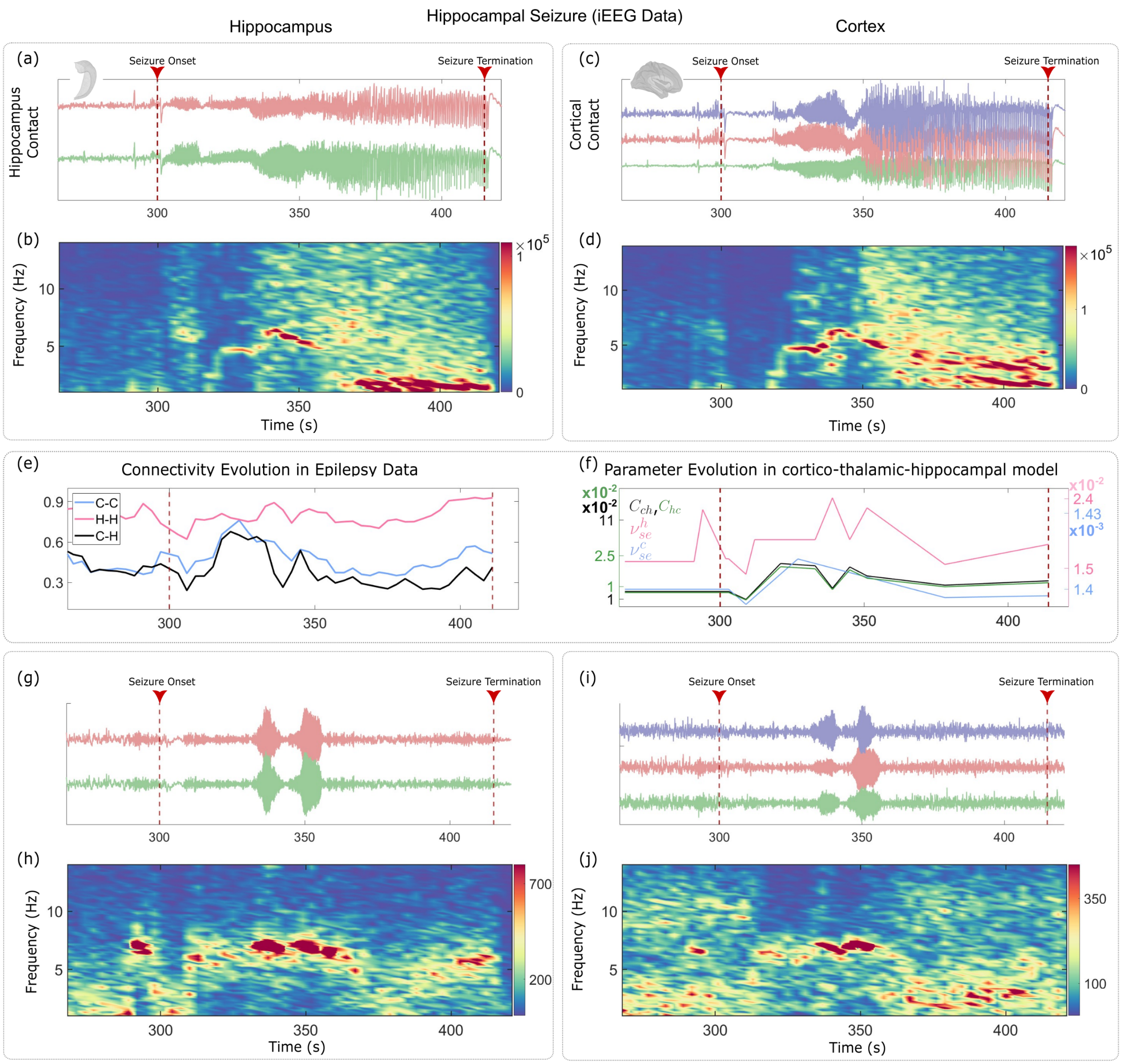


FIG. S6. Empirical and simulated seizure dynamics. (a) Preprocessed intracranial EEG (iEEG) traces from two hippocampal electrodes and (c) three cortical electrodes. Dotted red lines mark the onset and termination of the seizure as annotated by an epileptologist. (b,d) Average Short-time Fourier transform (STFT) spectrogram of the (b) hippocampal electrodes and (d) cortical electrodes. (e) Evolution of synchrony across unique hippocampal electrode pairs (pink), cortical electrode pairs (blue), and cortico–hippocampal electrode pairs (black). (f) System parameter trajectories inferred from the dynamic synchrony evolution in (e): hippocampal/cortical excitatory drive ($\nu^h_{se}/\nu^c_{se}$) and cortico-hippocampal coupling ($C_{ch}$ or $C_{hc}$). (g,i), Simulated time series from representative hippocampal (g) and cortical (i) vertices. (h, j), Simulated spectrogram from a representative hippocampal (h) and cortical vertex (j).

$$D_c(\mathbf{r},t)\phi^c_e(\mathbf{r},t) = \iint W_{ch}(\mathbf{r}-\mathbf{r}', t-t')\phi^h_e(\mathbf{r}',t')\, dt' d^2\mathbf{r}' + Q^c_e(\mathbf{r},t), \quad \text{(S7)}$$

$$D_h(\mathbf{r},t)\phi^h_e(\mathbf{r},t) = \iint W_{hc}(\mathbf{r}-\mathbf{r}', t-t')\phi^c_e(\mathbf{r}',t')\, dt' d^2\mathbf{r}' + Q^h_e(\mathbf{r},t). \quad \text{(S8)}$$

Note that the differential operators remain constant in form in this case. The coupling functions $W_{ch}$ and $W_{hc}$, which govern the interaction between the surfaces, are not point-to-point mappings. Instead, they are defined as proximity-based interactions that facilitate a smooth and continuous transport of fields between surfaces with potentially different spatial resolutions and geometries. The smooth and continuous transport of fields between the source and target surfaces is achieved by dynamically centering the spatial kernel $e^{-\kappa\|\mathbf{r}-\mathbf{r}'\|}$ on the source vertex that is closest to the target vertex $\mathbf{r}$. This ensures that the coupling is proximity-preserving, allowing each target vertex to receive a weighted contribution from source points in a spatially coherent manner. By integrating over the local neighborhood of the target vertex on the source surface, continuity is preserved and abrupt transitions are avoided. Equations S5 and S6 in the Fourier domain can be written as:

$$D_c(k,\omega)\phi_e^c(k,\omega) = W_{ch}(k,\omega)\phi_e^h(k,\omega) + Q_e^c(k,\omega), \quad \text{(S9)}$$

$$D_h(k,\omega)\phi_e^h(k,\omega) = W_{hc}(k,\omega)\phi_e^c(k,\omega) + Q_e^h(k,\omega). \quad \text{(S10)}$$

In the absence of external inputs and coupling between the sheets, the activity fields in the individual sheets satisfy the dispersion relations,

$$D_c(k,\omega)\phi_e^c(k,\omega) = 0, \quad \text{(S11)}$$

$$D_h(k,\omega)\phi_e^h(k,\omega) = 0, \quad \text{(S12)}$$

or, equivalently, $D_c(k,\omega) = 0$ and $D_h(k,\omega) = 0$. This indicates that, in isolation, the two neural sheets (cortex and hippocampus) exhibit intrinsic dynamics governed by their respective differential operators [9, 10]. However, when coupling is introduced, the combined dispersion relation is,

$$G(k,\omega) \equiv D_c(k,\omega)D_h(k,\omega) - W_{ch}(k,\omega)W_{hc}(k,\omega) = 0. \quad \text{(S13)}$$

Equation S13 represents the modified dispersion relation that governs how activity propagates under cortico-hippocampal coupling. If the coupling term $W_{ch}W_{hc}$ is zero, the cortical and hippocampal sheets will have resonant frequencies $\omega_c(k)$ and $\omega_h(k)$ respectively. If the product $W_{ch}W_{hc} \equiv \epsilon$ is small, perturbation theory can be applied to obtain corrections to $\omega_c(k)$ and $\omega_h(k)$ up to $\mathcal{O}(\epsilon)$. The zeros of the function $G(k,\omega)$ can be written as $\omega_c^* = \omega_c + \delta\omega_c$ and $\omega_h^* = \omega_h + \delta\omega_h$.

Expanding $G(k,\omega_c+\delta\omega_c)$ around $\omega_c$ gives,

$$\begin{aligned} G(k,\omega_c+\delta\omega_c) = {} & D_c(k,\omega_c)D_h(k,\omega_c) - \epsilon \\ & + D_c'(k,\omega_c)D_h(k,\omega_c)\,\delta\omega_c \\ & + \mathcal{O}(\epsilon^2), \end{aligned} \quad \text{(S14)}$$

where $D' = \partial D(k,\omega)/\partial\omega$. Because $\omega_c$ is the root of the uncoupled equation $D_c(k,\omega) = 0$, equation S14 simplifies to:

$$G(k,\omega_c+\delta\omega_c) = -\epsilon + D_c'(k,\omega_c)D_h(k,\omega_c)\delta\omega_c + \mathcal{O}(\epsilon^2). \quad \text{(S15)}$$

Now, $G(k,\omega_c+\delta\omega_c) = 0$, given the assumption of perturbed roots of the coupled function. Therefore, S15 can be rearranged to,

$$\delta\omega_C = \frac{\epsilon}{D_C'(k,\omega_c)D_H(k,\omega_c)}. \quad \text{(S16)}$$

Performing a similar expansion around $\omega_h$ gives:

$$\delta\omega_H = \frac{\epsilon}{D_H'(k,\omega_h)D_C(k,\omega_h)}. \quad \text{(S17)}$$

This shows that as the cortico-hippocampal coupling is gradually introduced, a shift in the resonant frequencies will arise as a natural consequence of the coupling, providing a theoretical basis for the frequency changes observed in the numerical simulations as the coupling strength is increased.

#### S2.3. Modeling the Coupling Functions

The coupling functions $W_{ch}$ and $W_{hc}$ must be expressed in a continuous space. The coupling strength between the cortical and hippocampal maps in the present manuscript is modeled using exponentially decaying functions. The spatial part of the coupling functions are defined as,

$$w_{ch}(\mathbf{r},\mathbf{r}') = \begin{cases} e^{-\kappa\|\mathbf{r}-\mathbf{r}'\|}, & \text{if } \mathbf{r}' \in \mathcal{N}_{99}(r_s), \\ 0, & \text{otherwise,} \end{cases} \quad \text{(S18)}$$

$$w_{hc}(\mathbf{r},\mathbf{r}') = \begin{cases} e^{-\kappa\|\mathbf{r}-\mathbf{r}'\|}, & \text{if } \mathbf{r}' \in \mathcal{N}_{99}(r_s), \\ 0, & \text{otherwise.} \end{cases} \quad \text{(S19)}$$

where $r_s = \arg\min_{\mathbf{r}'\in\mathcal{M}_S} \|\mathbf{r}-\mathbf{r}'\|$ is the nearest source vertex to the target vertex $\mathbf{r}$, and $\mathcal{N}_{99}(r_s)$ denotes its 99 nearest neighboring vertices on the mapped domain.

By defining the coupling adaptively, the spatial kernel remains proximity-preserving and avoids distortions from the mapping. Since the coupling functions $w_{ch}(\mathbf{r},\mathbf{r}')$ and $w_{hc}(\mathbf{r},\mathbf{r}')$ decay exponentially with distance, the nearest-neighbor cutoff effectively limits interactions to a finite range. Beyond this range, contributions rapidly approach zero, making the cutoff a practical approximation of infinite-range decay. Hence, Eqs. S18 and S19 simplify to,

$$w_{ch}(\mathbf{r}-\mathbf{r}') = e^{-\kappa\|\mathbf{r}-\mathbf{r}'\|}, \quad \text{(S20)}$$

$$w_{hc}(\mathbf{r}-\mathbf{r}') = e^{-\kappa\|\mathbf{r}-\mathbf{r}'\|}. \quad \text{(S21)}$$

Given a constant time delay of $\tau_{ch}$ for the field transport from the cortical surface to the hippocampal surface and vice versa, we can separate the temporal and spatial parts of the coupling function and represent the temporal coupling part with a Dirac-delta function. This modifies Eq S7 and S8 to,

$$D_c(\mathbf{r},t)\,\phi_e^c(\mathbf{r},t) = \int_{\mathbb{R}^2}\int_{-\infty}^{\infty} C_{ch}w_{ch}(\mathbf{r}-\mathbf{r}')\,\delta(t-t'-\tau_{ch}) \times \phi_e^h(\mathbf{r}',t')\,dt'\,d^2\mathbf{r}' + Q_e^c(\mathbf{r},t), \quad \text{(S22)}$$

$$D_h(\mathbf{r},t)\,\phi_e^h(\mathbf{r},t) = \int_{\mathbb{R}^2}\int_{-\infty}^{\infty} C_{hc}w_{hc}(\mathbf{r}-\mathbf{r}')\,\delta(t-t'-\tau_{ch}) \times \phi_e^c(\mathbf{r}',t')\,dt'\,d^2\mathbf{r}' + Q_e^h(\mathbf{r},t). \quad \text{(S23)}$$

Since the coupling functions $w_{ch}(\mathbf{r}-\mathbf{r}')$ and $w_{hc}(\mathbf{r}-\mathbf{r}')$ have the same functional form, and the differential operators $D_c$ and $D_h$ are of the same structure, the equations governing the cortical and hippocampal activity fields take an identical form. Thus, without loss of generality, analysis can be done with only one of these equations.

The Dirac delta function simplifies the temporal convolution integral as follows,

$$\int_{-\infty}^{\infty} \delta(t-t'-\tau_{ch})f(t')\,dt' = f(t-\tau_{ch}). \quad \text{(S24)}$$

Applying this property to the coupling term in the first half of Equation S22 gives,

$$\int_{-\infty}^{\infty} \delta(t-t'-\tau_{ch})\phi_e^h(\mathbf{r}',t')\,dt' = \phi_e^h(\mathbf{r}',t-\tau_{ch}). \quad \text{(S25)}$$

Therefore, the cortical field equation S22 simplifies to,

$$D_c(\mathbf{r},t)\phi_e^c(\mathbf{r},t) = C_{ch}\int_{\mathbb{R}^2} w_{ch}(\mathbf{r}-\mathbf{r}')\phi_e^h(\mathbf{r}',t-\tau_{ch})\,d^2\mathbf{r}' + Q_e^c(\mathbf{r},t). \quad \text{(S26)}$$

The Fourier transform of a function $f(\mathbf{r},t)$ can be defined as,

$$F(k,\omega) = \int_{\mathbb{R}^2}\int_{-\infty}^{\infty} f(\mathbf{r},t)e^{-i(k\cdot\mathbf{r}-\omega t)}\,dt\,d^2\mathbf{r}. \quad \text{(S27)}$$

The functional form for the cortico-hippocampal field coupling is,

$$W_{ch}(\mathbf{r},t) = \int_{\mathbb{R}^2} w_{ch}(\mathbf{r}-\mathbf{r}')\phi_e^h(\mathbf{r}',t-\tau_{ch})\,d^2\mathbf{r}'. \quad \text{(S28)}$$

Taking a Fourier transform gives,

$$W_{ch}(k,\omega) = \int_0^{\infty}\int_{\mathbb{R}^2}\left[\int_{\mathbb{R}^2} w_{ch}(\mathbf{r}-\mathbf{r}')\phi_e^h(\mathbf{r}',t-\tau_{ch})d^2\mathbf{r}'\right] e^{-i(k.\mathbf{r}-\omega t)}\,d^2\mathbf{r}\,dt, \quad \text{(S29)}$$

where, $\int_{\mathbb{R}^2} w_{ch}(\mathbf{r}-\mathbf{r}')\phi_e^h(\mathbf{r}',t-\tau_{ch})d^2\mathbf{r}'$ represents the 2-D convolution of the coupling function $w_{ch}(.)$ with $\phi_e^h(\mathbf{r},t-\tau_{ch})$.

By performing the spatial integral first,

$$W_{ch}(k,\omega) = \int_0^{\infty}\left[\int_{\mathbb{R}^2} w_{ch}(\mathbf{r}) * \phi_e^h(\mathbf{r},t-\tau_{ch})e^{-ik.\mathbf{r}}d^2\mathbf{r}\right] e^{i\omega t}dt, \quad \text{(S30)}$$

$$W_{ch}(k,\omega) = \int_0^{\infty}\left[w_{ch}(k)\phi_e^h(k,t-\tau_{ch})\right] e^{i\omega t}dt, \quad \text{(S31)}$$

$$W_{ch}(k,\omega) = w_{ch}(k)e^{i\omega\tau_{ch}}\phi_e^h(k,\omega). \quad \text{(S32)}$$

Focusing on the functional form of the spatial part of the coupling kernel: $w_{ch}(\mathbf{r}) = e^{-\kappa\mathbf{r}}$,

$$w_{ch}(k) = \int e^{-\kappa\mathbf{r}}e^{-ik\cdot\mathbf{r}}\,d^2\mathbf{r}. \quad \text{(S33)}$$

Using polar coordinates,

$$w_{ch}(k) = \int_0^{2\pi}\int_0^{\infty} e^{-\kappa\mathbf{r}}e^{-ikr\cos(\theta)}\mathbf{r}\,d\mathbf{r}\,d\theta. \quad \text{(S34)}$$

The angular integral in the above equation gives a Bessel function of the first kind,

$$\int_0^{2\pi} e^{-ik.\mathbf{r}\cos(\theta)}\,d\theta = 2\pi J_0(k.\mathbf{r}). \quad \text{(S35)}$$

Thus, the Fourier transform becomes,

$$w_{ch}(k) = 2\pi\int_0^{\infty} e^{-\kappa r}J_0(k.\mathbf{r})\mathbf{r}\,d\mathbf{r}, \quad \text{(S36)}$$

$$w_{ch}(k) = \frac{2\pi\kappa}{(\kappa^2+k^2)^{3/2}}, \quad \text{(S37)}$$

where $k$ is the spatial wavevector and $\kappa$ is the exponent for the spatial decay of the coupling. Hence, equation S9 for the cortical surface becomes,

$$D_c(k,\omega)\phi_e^c(k,\omega) = Q_e^c(k,\omega) + C_{ch}w_{ch}(k)\cdot e^{i\omega\tau_{ch}}\phi_e^h(k,\omega), \quad \text{(S38)}$$

which indicates that, in the idealized setting of a continuous and translation-invariant domain, each wavevector $k$ of the hippocampal field is coupled directly to the same $k$ in the cortical field, with an amplitude $w_{ch}(k)$ determined by the Fourier transform of the coupling kernel. However, the finite geometry and curved topology of the cortical and hippocampal surfaces replace the continuous $k$-space with discrete Laplace-Beltrami eigenmodes that are determined by the surfaces' intrinsic geometry and boundary conditions. Hence, the numerical scheme implemented in this work, which employs an eigenmode basis, serves as an approximation to the continuous $k$-space framework. Although strict one-to-one coupling between individual modes is not guaranteed in such geometries, the radially symmetric and exponentially decaying nature of the coupling function tends to preserve spectral locality. As a result, interactions are predominantly facilitated between modes of comparable characteristic wavelengths, supporting the intuition that the cortico-hippocampal coupling preferentially links spatial patterns of similar scales across the two neural sheets.

#### S2.4. Mode-Mode coupling in the Corticothalamic/hippocampo-septal System

The nonlinear feedback loops introduced by the sigmoid function in the corticothalamic/hippocampo-septal systems can lead to mode-mode coupling between different cortical or hippocampal modes. These nonlinearities and feedback mechanisms can cause interactions between different eigenmodes of the system, leading to complex dynamics that are not present in linear systems. In this section, a mathematical heuristic is used to demonstrate the existence of mode-mode coupling in the corticothalamic or the hippocampo-septal loop. The neural field equation for the field activity on either surface $\phi_e^a(\mathbf{r},t)$ is,

$$\left[\frac{1}{\gamma_a^2}\frac{\partial^2}{\partial t^2}+\frac{2}{\gamma_a}\frac{\partial}{\partial t}+1-r_a^2\nabla^2\right]\phi_e^a(\mathbf{r},t)=Q_e^a(\mathbf{r},t). \quad \text{(S39)}$$

The field term ($\phi_e^a(\mathbf{r},t)$) and source term ($Q_e^a(\mathbf{r},t)$) can be expanded in terms of the eigenmodes $R_n^a(\mathbf{r})$ of the Laplacian on the cortical or the hippocampal surface,

$$\phi_e^a(\mathbf{r},t)=\sum_{n=1}^{N_a}\zeta_a^n(t)R_n^a(\mathbf{r}), \quad \text{(S40)}$$

$$Q_e^a(\mathbf{r},t)=\sum_{n=1}^{N_a}\rho_n^a(t)R_n^a(\mathbf{r}). \quad \text{(S41)}$$

The eigenmodes satisfy the Helmholtz equation,

$$\nabla^2R_n^a(\mathbf{r})=-\lambda_n^aR_n^a(\mathbf{r}). \quad \text{(S42)}$$

Substituting this into the wave equation, the governing equation for each eigenmode $n$ becomes,

$$\left[\frac{1}{\gamma_a^2}\frac{d^2}{dt^2}+\frac{2}{\gamma_a}\frac{d}{dt}+1+r_a^2\lambda_n^a\right]\zeta_n^a(t)=\kappa_n^a(t), \quad \text{(S43)}$$

which represents a set of uncoupled linear ODEs for each eigenmode. In a linear system, where $Q_e^a\propto V_e^a$, all the eigenmodes evolve independently. However, the source term $Q_e^a(r,t)$ for the wave equation depends non-linearly on the transmembrane potential $V_a(r,t)$ via the sigmoid function as follows,

$$Q_j^a(\mathbf{r},t)=\frac{(Q_j^a)^{\max}}{1+\exp\left(-\frac{V_j^a(\mathbf{r},t)-\theta_j^a}{\sigma_j^a}\right)} \quad \text{(S44)}$$

where, $j$ represents the neuronal population and $Q_e^a(\mathbf{r},t)$ represents the firing rate for the excitatory neuronal population on the surface $a$ ($a=c$ for cortex and $a=h$ for hippocampus). The transmembrane potential $V_j^a(\mathbf{r},t)$ depends on $P_j^a(\mathbf{r},t)$, which aggregates inputs from other neuronal populations via the linear differential equation

$$\left[\frac{1}{\alpha_a\beta_a}\frac{d^2}{dt^2}+\left(\frac{1}{\alpha_a}+\frac{1}{\beta_a}\right)\frac{d}{dt}+1\right]V_j^a(\mathbf{r},t)=P_j^a(\mathbf{r},t), \quad \text{(S45)}$$

where, $P_j^a(\mathbf{r},t)$ represents the sum of inputs to the neural population $j$ on the cortical ($a=c$) or the hippocampal ($a=h$) surface and $j$ represents the neural populations (the relay or the reticular nuclei of the thalamus or the septum or the excitatory/inhibitory neural population on the cortical or the hippocampal surfaces).

Moreover, $P_j^a(\mathbf{r},t)\ \forall j=r,s,e$, can be written as follows,

$$\begin{aligned}P_e^a(\mathbf{r},t)&=\nu_{ee}^a\phi_e^a(\mathbf{r},t)+\nu_{ei}^a\phi_i^a(\mathbf{r},t)+\nu_{es}^aQ_s^a(\mathbf{r},t-\tau_a)\\P_r^a(\mathbf{r},t)&=\nu_{rs}^aQ_s^a(\mathbf{r},t)+\nu_{re}^a\phi_e^a(\mathbf{r},t-\tau_a)\\P_s^a(\mathbf{r},t)&=\nu_{se}^a\phi_e^a(\mathbf{r},t-\tau_a)+\nu_{sr}^aQ_r^a(\mathbf{r},t)+\nu_{sn}^a(\phi_{n0}+\eta_a),\end{aligned} \quad \text{(S46)}$$

where, $\phi_e^a$ represents the excitatory ($e$) field on the cortical/hippocampal ($a$) surface. Even though the ODE for $V_j^a$ is linear, the presence of $Q_j^a$ (which is a nonlinear function of $V_j^a$) in the aggregated input $P_j^a$ introduces nonlinear dependencies.

To summarize, the governing equations form a closed loop ($Q_j^a\to\phi_e^a\to V_j^a\to Q_j^a$) with nonlinear dependencies.

- Each $Q_j^a$ (e.g., $Q_s^a, Q_r^a$) is a sigmoid function of its respective $V_j^a$.
- These $Q_j^a$ terms appear in the input $P_j^a$ for the equations governing the evolution of the transmembrane potential $V_j^a$ for other populations (Equations S46 and S45).
- Thus, nonlinearities in $Q_j^a$ all feed back into the equations for $V_j^a$, even though the ODE for $V_j^a$ is linear.

Assuming that the system operates near a fixed point $V_{j0}^a$, The sigmoid function $Q_j^a(V_j^a)$ can be expanded about $V_{j0}^a$ to third order as follows,

$$Q_j^a \approx Q_{j0}^a + \left.\frac{dQ_j^a}{dV_j^a}\right|_{V_{j0}^a} (V_j^a - V_{j0}^a) + \frac{1}{2}\left.\frac{d^2Q_j^a}{d(V_j^a)^2}\right|_{V_{j0}^a} (V_j^a - V_{j0}^a)^2 + \frac{1}{6}\left.\frac{d^3Q_j^a}{d(V_j^a)^3}\right|_{V_{j0}^a} (V_j^a - V_{j0}^a)^3. \quad \text{(S47)}$$

The cubic term is critical for mode coupling. Defining $\delta V_j^a = V_j^a - V_{j0}^a$, the cubic term can be written as,

$$\frac{1}{6}\left.\frac{d^3Q_j^a}{d(V_j^a)^3}\right|_{V_{j0}^a} \delta V_j^{a3}. \quad \text{(S48)}$$

Because the eigenmodes form a complete orthonormal basis on the cortical/hippocampal surface, $\delta V_j^a(\mathbf{r}, t)$ can be expanded as,

$$\delta V_j^a(\mathbf{r}, t) = \sum_{n=1}^{N_a} \rho_n^a(t) R_n^a(\mathbf{r}). \quad \text{(S49)}$$

Cubing $\delta V_j^a$ gives,

$$\delta V_j^{a3}(\mathbf{r}, t) = \left(\sum_{n=1}^{N_a} \rho_n^a(t) R_n^a(\mathbf{r})\right)^3 = \sum_{n,m,k} \rho_n^a(t)\rho_m^a(t)\rho_k^a(t) R_n^a(\mathbf{r}) R_m^a(\mathbf{r}) R_k^a(\mathbf{r}). \quad \text{(S50)}$$

These products of eigenmodes are projected back onto the eigenmode basis,

$$\mu_p^a(t) = \sum_{n,m,k} \rho_n^a(t)\rho_m^a(t)\rho_k^a(t) C_{nmkp}^a, \quad \text{(S51)}$$

where,

$$C_{nmkp}^a = \int_{\mathcal{M}_a} R_n^a(\mathbf{r}) R_m^a(\mathbf{r}) R_k^a(\mathbf{r}) R_p^a(\mathbf{r})\, dS. \quad \text{(S52)}$$

The coefficients $C_{nmkp}^a$ quantify the degree to which the product $R_n^a R_m^a R_k^a$ projects onto $R_p^a$.

As an example, suppose,

$$\delta V_j^a(\mathbf{r}, t) = \rho_1^a R_1^a(\mathbf{r}) + \rho_2^a R_2^a(\mathbf{r}), \quad \text{(S53)}$$

then,

$$\delta V_j^{a3} = \rho_1^{a3} R_1^{a3} + 3\rho_1^{a2}\rho_2^a R_1^{a2} R_2^a + 3\rho_1^a \rho_2^{a2} R_1^a R_2^{a2} + \rho_2^{a3} R_2^{a3}. \quad \text{(S54)}$$

Projecting $R_1^{a2} R_2^a$ onto $R_p^a$ yields,

$$C_{112p}^a = \int_{\mathcal{M}_a} R_1^{a2} R_2^a R_p^a\, dS. \quad \text{(S55)}$$

A nonzero $C_{112p}^a$ indicates that mode $p$ is driven by a term proportional to $\rho_1^{a2}\rho_2^a$, demonstrating mode coupling.

Hence, the cubic nonlinearity in the sigmoid is shown to induce interactions between modes through these coupling coefficients, which are absent in linear systems.

[9] P. Robinson, P. Loxley, S. O’connor, and C. Rennie. Modal analysis of corticothalamic dynamics, electroen-

cephalographic spectra, and evoked potentials. *Physical Review E*, 63(4):041909, 2001.

[10] P. A. Robinson, C. J. Rennie, and D. L. Rowe. Dynamics of large-scale brain activity in normal arousal states and epileptic seizures. *Physical Review E*, 65(4):041924, 2002.
